# Backwards compatibility to classical experiments grounds beta responses to naturalistic speech in temporal acoustic forecasting

**DOI:** 10.64898/2026.03.18.712602

**Authors:** Christoph Daube, Joachim Gross, Robin A. A. Ince

## Abstract

Current neuroscience is shifting from simple controlled paradigms towards rich and ecologically valid naturalistic stimuli. Correspondingly, insights from historic “impoverished” artificial paradigms are considered to be seriously challenged by generalisation to modern naturalistic contexts. Here, we argue that the reverse, backwards compatibility, is an under-appreciated and easily accessible benchmark and show that it disambiguates model comparison beyond naturalistic stimuli. We analyse magnetoencephalography (MEG) beta power responses to an audiobook stimulus using canonical correlation analysis (CCA) and replicate reported links between beta power and syntactic parsing. However, a simple stimulus-computable acoustic model predicts the same variance, suggesting a domain-general rather than linguistic function of beta responses. We therefore test the backwards compatibility of speech-trained models to classic rhythmic tones. While generalisation initially fails, reducing hidden and unnecessary degrees of freedom of the models’ phase responses allows successful generalisation. Crucially, this greatly improves model adjudication: Several models that perform indistinguishably on speech differ in how well they predict responses to simple tones. In this comparison, a simple forecasting deep neural network (DNN) outperforms acoustics by internalising a “slow-decay” prior as a structural mirror of sluggish speech dynamics. This grounds beta responses in canonical temporal forecasting, bridging modern naturalism to established auditory psychophysics.

## Introduction

How the human brain derives meaning from speech is a central question in neuroscience. In trying to solve this puzzle, research has developed from early approaches rooted in psychophysics and controlled simplistic stimulus material all the way to modern naturalistic neuroimaging using rich and ecologically valid speech sounds. The latter allow us to observe the system under study in ubiquitous settings typical of its natural operation, rather than in unfamiliar ones, and thus promise to elicit vigorous neural responses (David et al., 2004; Rieke et al., 1995; Theunissen et al., 2000). This is usually considered to reflect scientific progress (Hamilton & Huth, 2020), reducing older and simpler approaches to mere “baby steps” that lack a “big picture” (VanRullen, 2017), and casting modern naturalistic approaches as the logical consequence of increasing methodological sophistication: Contemporary machine learning allows to extract ever more informative features directly from complex waveforms of environmental sounds, music or speech, and these features can be used to build “stimulus-computable” (Giordano et al., 2023; Kell et al., 2018; Kriegeskorte & Douglas, 2018) models of brain responses to such stimuli (Naselaris et al., 2011).

This is thought to be out of reach for models developed from more simplistic scenarios, hampering their generalisation to naturalistic settings. Conversely, it seems tempting to assume that once a model captures the high-dimensional variance of complex naturalistic data, its performance on simplistic laboratory stimuli becomes trivial. It is thus not surprising that a substantial amount of research documenting models of neural speech processing is limited to evaluating predictive performance of speech-trained models *within-distribution*, that is, on speech data (Brodbeck et al., 2025; Daube et al., 2019; Di Liberto et al., 2015; Ding & Simon, 2012; Donhauser & Baillet, 2020; Heilbron et al., 2022; O’Sullivan et al., 2015).

It is however increasingly (re-)appreciated that a multitude of models with very different properties can achieve indistinguishably high performance when evaluated on uncontrolled data sets (Conwell et al., 2024; Daube et al., 2019; Golan et al., 2020, 2022). In other words, if models are our hypotheses of brain function, then testing their predictive performance on naturalistic data does not allow us to accept or reject them. This form of *multiple realisability* (Edelman & Gally, 2001; Putnam, 1967) thus makes it hard to adjudicate between different models and in this way impedes scientific progress.

Further, it has been pointed out that simply chasing for prediction performance on uncontrolled datasets leaves it unclear what specific features in fact drive those predictions (Bowers et al., 2022; Schyns et al., 2022). Controlled experiments in turn can test specific hypotheses by manipulating corresponding stimulus factors while holding others constant (Ding et al., 2016; Golan et al., 2022; Rust & Movshon, 2005; Schyns et al., 2023). Intriguingly, models that succeed on naturalistic stimuli have been shown to struggle with generalising to classic controlled experiments known from the literature (Bowers et al., 2022). The field’s history is thus not at all eclipsed by its contemporary practice.

Here, we thus suggest harnessing the knowledge we already have at our fingertips in the form of documented controlled experiments to disambiguate model adjudication beyond what is possible with naturalistic stimulus material. We contend that testing models with regards to such *backwards compatibility* (Figure 1) is an under-appreciated benchmark that can easily enrich any model comparison limited to naturalistic test cases with more diagnostic *out-of-distribution* stimuli. It can further point to overlooked computational motives worth considering when devising candidate models. Eventually, these should thrive in both naturalistic and controlled settings (Carvalho & Lampinen, 2025; Schrimpf et al., 2020).

**Figure 1:**
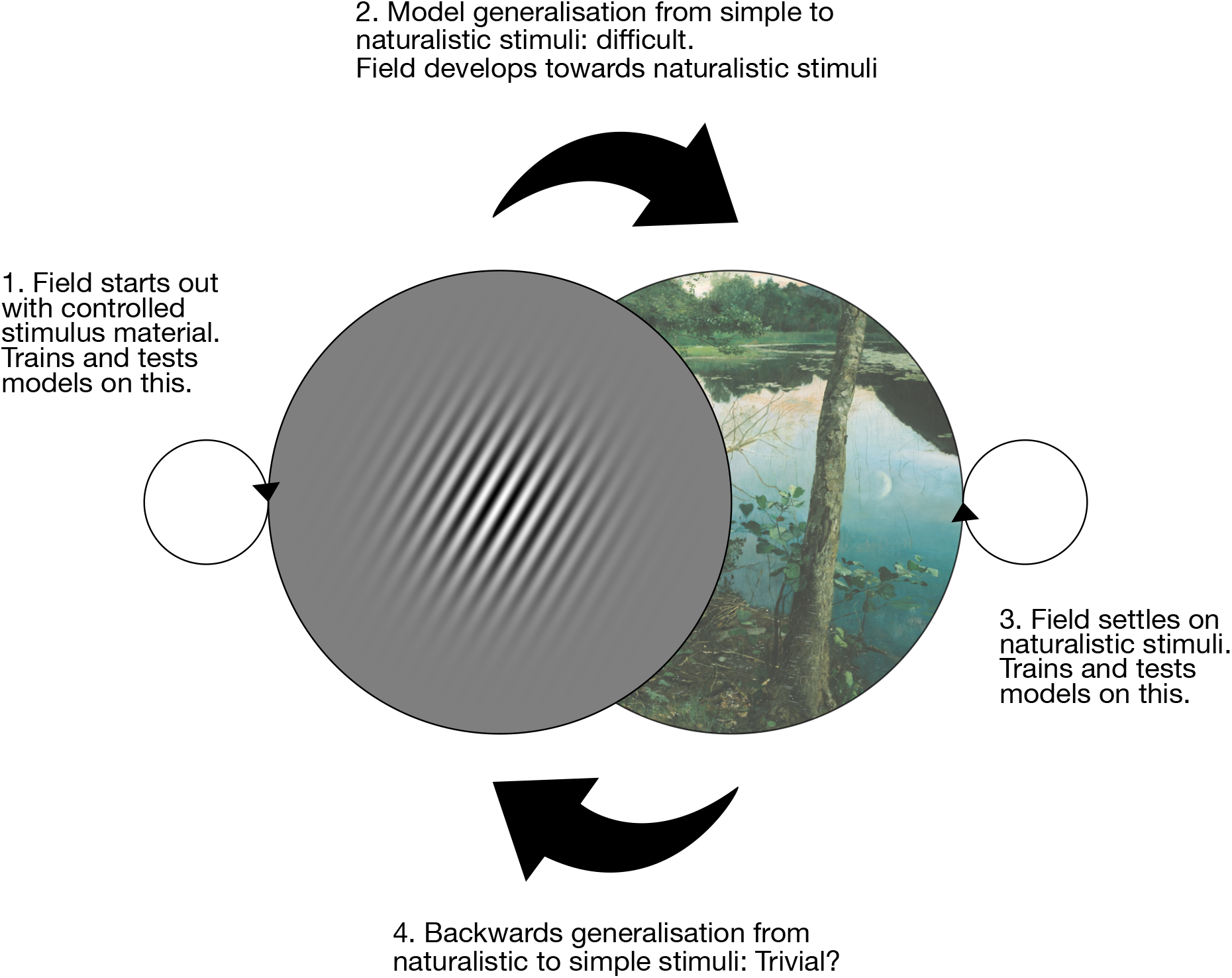
Backwards compatibility of encoding models. The transition from using simplistic controlled stimuli (symbolic Gabor patch, left) to naturalistic stimuli (right) is often viewed as a one-way path towards ecological validity. Here, we argue for the inclusion of the reverse path (bottom arrow): evaluating models based on their ability to recapitulate findings from classical controlled experiments. We propose that this “backwards compatibility” is an easily accessible out-of-distribution benchmark to improve adjudication between models trained on naturalistic data with similar within-distribution performance. Further, it can inspire computational motives for models that actually manage to account for variance in both settings. Image credit: Eilif Petersen, Summer Night.

To illustrate our point, we consider an emerging frontier in neuroimaging recorded at high temporal resolution during passive speech listening. When constructing encoding models of such responses, researchers usually focus on the clearly discernible neural time domain signal at frequencies slower than ∼10 Hz (Brodbeck & Simon, 2020; Crosse et al., 2016). Only recently, researchers have started to construct models of power time courses in the faster beta band (13 - 30 Hz). The results suggest that beta dynamics track high-level computations like syntactic parsing and statistical prediction, promising to offer a non-invasive window into the brain’s internal language model (e.g. Armeni et al., 2019; Weissbart & Martin, 2024; Zioga et al., 2023). However, it is difficult to conclusively rule out simpler explanations of such findings (Daube et al., 2019). It is for example possible that compressive nonlinearities of time varying sound energy exist which can boost performance beyond control models in similar ways as linguistic features. If this were the case, it would suggest that beta power during speech listening might in fact be reflective of a more fundamental domain-general auditory process rather than specific to linguistic operations. It should then be possible for models trained on speech to be backwards compatible to simple auditory paradigms: Work using non-linguistic stimuli has previously linked beta oscillations to temporal forecasting, i.e. the anticipation of upcoming sensory events based on rhythm and timing (Arnal et al., 2015; Betti et al., 2021; de Lange et al., 2013; Fujioka et al., 2012; Grabenhorst et al., 2025; Sedley et al., 2016).

We first develop a data-driven method to extract beta power dynamics from MEG data with high sensitivity. This allows us to replicate reports of the relevance of linguistic features, but also reveals competitive acoustic features. We therefore consider a domain general auditory functional significance of the beta signal and test the backwards compatibility of our models to a controlled rhythmic tones experiment known from the literature (Fujioka et al., 2012). We find that this generalisation is surprisingly non-trivial, and identify underappreciated and superfluous degrees of freedom in the linear encoding models. Constraining these timing-rather than feature-related degrees of freedom allows speech-trained models to generalise to the simple tone stimuli. Inspired by this generalisation to a task linked to temporal forecasting, we pit several stimulus-computable DNNs concerned with temporal forecasting against each other. While these perform highly similarly when tested on held-out speech data, backwards compatibility testing on tone data reveals strong differences between the models. In this way, we find that a canonical sound energy forecasting model best accounts for beta bursting dynamics in the human auditory cortex across naturalistic and artificial stimuli. Lastly, we show that its advantage over our best acoustic models is rooted in a “slow-decay” prior learned from speech data.

## Results

We re-analysed a dataset of 24 participants who had listened to an audiobook of 1 hour duration while their MEG had been recorded (Daube et al., 2019).

### Extracting power responses to speech using Canonical Correlation Analysis (CCA)

In the first step, we were interested in assessing if there was a systematic linear relationship between the speech stimulus and neural power dynamics at any frequency and at any sensor. To test this, we submitted the MEG data and a speech stimulus feature to a regularised CCA (Bilenko & Gallant, 2016; de Cheveigné et al., 2018; Stratos, 2020 Figure 1A), which we computed in a nested 6-fold cross-validation. In this way, we found linear weights to project the MEG sensor power time courses across a range of frequency bands (20 bands of 3Hz width from 1 to 40 Hz) onto a subspace that was maximally correlated with a linear combination of the time-lagged speech envelope.

The first component identified by this CCA yielded a correlation of stimulus and response projections of .2220 (mean across participants and folds), which decreased sharply for the following components (component 2: .0230, component 3: .0097, Figure 2B). We thus focused our analyses on the first component and did not further consider the remaining components. Importantly, our top component clearly outperformed encoding models trained to predict individual grid points from a classic beamformer inverse solution: When taking the peak of encoding model performances in each individual participant’s whole brain map, this averaged to a correlation of .1868. When comparing the two approaches with a Bayesian approach, a fraction *f*_*h*_ *=* 0.9989 of the posterior samples indicated “decisive” (Jeffreys, 1998) evidence for CCA providing higher sensitivity.

**Figure 2:**
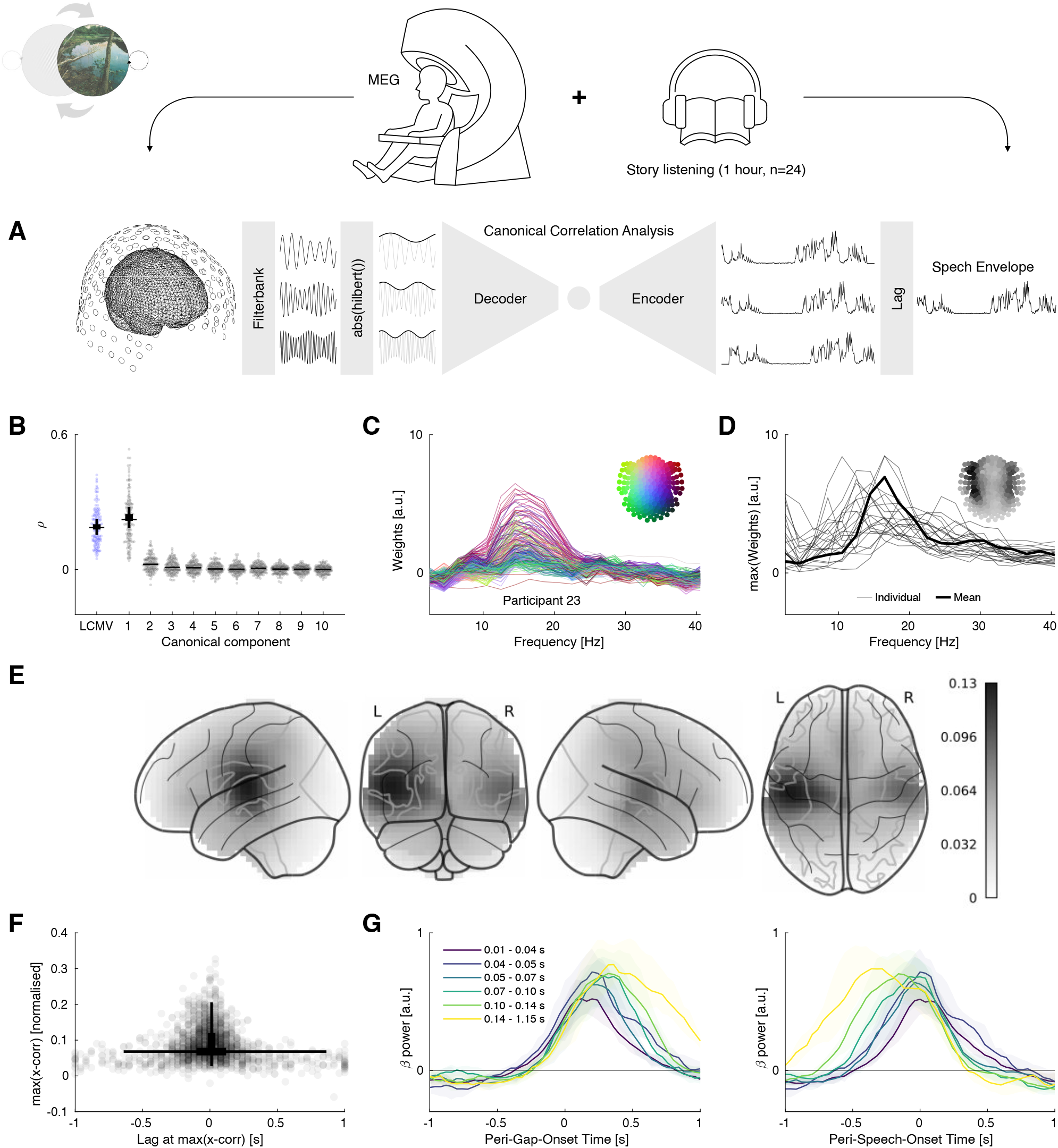
Extracting neural power dynamics in response to speech with CCA. **A**) We prepare MEG power time courses at a range of frequencies from sensor level MEG data recorded during passive audiobook listening. We also prepare the lagged stimulus envelope and fit a regularised CCA between stimulus and response variables. This results in linear “decoders” mapping neural responses onto canonical response components and linear “encoders” mapping the stimulus feature onto canonical stimulus components that are maximally correlated with the canonical response components. **B**) The first canonical component achieves the highest canonical correlation, further components only achieve negligible levels This yields a stronger signal than the peak grid point of a linearly constrained minimum variance (LCMV, blue) inverse solution. **C**) Weights of the decoder (Haufe et al., 2014) from an exemplary participant, resolved over frequencies and sensors. Inset maps colours to sensor positions.Weights point to right-lateralised temporal sensors and a frequency range between 10 and 20 Hz contributing to the decoded response component. **D**) Maximum (over sensors) of decoder weights at each frequency for each of n=24 participants. Inset shows average sensor topographies. Thin lines show individual participants, bold line shows average across participants. **E**) Grand average source localisation using linear model to predict first canonical response component from beamformer power time series estimates of the whole brain. Source plots report squared Pearson correlation between test-set predictions and first canonical response components. **F**) Upper triangular of all between-participant cross-correlations (maximum value over lags) of canonical response components. Most cross-correlations peak at short latencies. **G**) Canonical response components epoched according to gap onsets (left) and speech onsets (right), grand average. Colour codes different gap lengths. Gap length correlates with bursting duration.

We multiplied the spatio-spectral MEG response filters of the CCA (“decoders”) with the MEG sensor covariance matrices to obtain interpretable pattern matrices (Haufe et al., 2014). These showed that bilateral temporal sensors in the beta frequency band (13 − 30 Hz) contributed most to the canonical response components (Figure 2C, D). To source-localise these components, we estimated cross-validated linear models (Parra et al., 2005) that predicted the canonical response components from power time series at individual source locations as estimated by a linearly constrained minimum variance beamformer (Van Veen et al., 1997). The resulting source maps of test performance suggested bilateral superior temporal cortices (Figure 2E) as the primary generators of our canonical response components. We were next interested in the consistency across subjects of these response projections. We cross-correlated the canonical response components for each of the 276 participant pairs (Figure 2F), which peaked at a delay of -.0094s ± .1479 (mean ±SD, standard deviation), and at these peaks, the correlation was (mean ±SD) .0807 ± .0482. While some pairs of participants were most similar at relatively long latencies, the corresponding peaks had relatively low correlation values and the bulk of pairs was most correlated at short latencies (95% shorter than ± .28 s), suggesting similar temporal dynamics across participants. Given previous findings in the beta band (Fujioka et al., 2012), we hypothesised that the correlation between response projections and the time-lagged envelope was driven by high beta power values (“bursts”) preceding speech onsets. We therefore epoched the response projections around stimulus energy increases following gaps, and found that the response projections indeed peaked close to these onsets (Figure 2G). This was highly similar regardless of whether the epoching was with respect to the onsets of the gaps or the onsets of speech following the gaps, and in both cases, we observed stronger and more sustained bursting for longer gaps.

Taken together, we developed an approach to robustly extract neuronal power responses to continuous passive speech listening. We found this to provide a stronger signal than an off-the-shelf beamformer inverse solution. This revealed a response component in the beta band, which we source-localised to superior temporal cortices. A preliminary analysis of this response component showed that it increases during short gaps in the speech stimulus.

### Replication of high-level effects and explanation in terms of acoustic features

In a second step, we were interested in comparing more detailed models of the identified canonical response component. To do so, we transformed the stimulus material into a range of “linearising feature spaces” (Naselaris et al., 2011 Figure 2B). We then linearly re-predicted the same canonical response components we reported in the previous section from this range of feature spaces (Figure 3A) and compared the Pearson correlations between observations and predictions.

**Figure 3:**
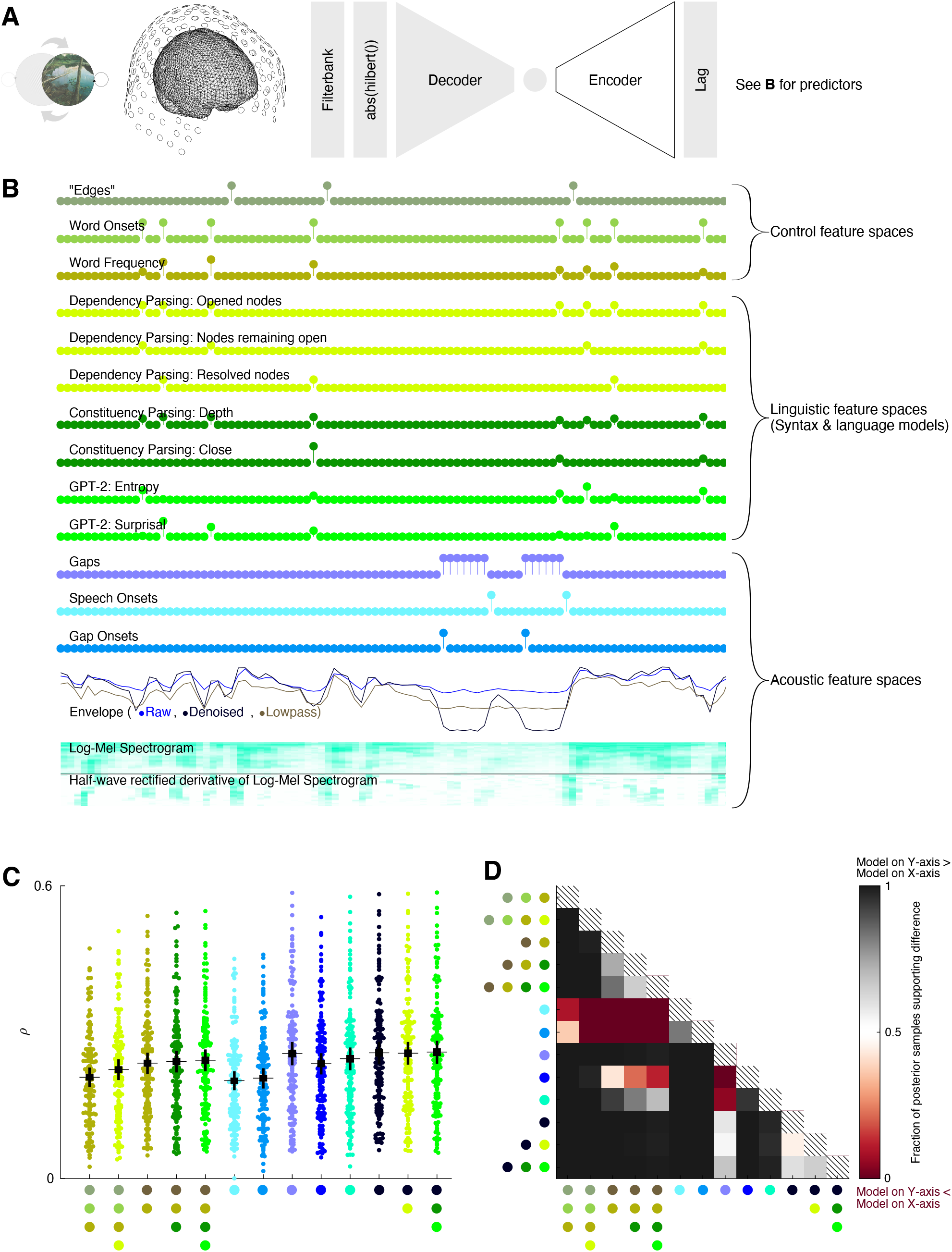
Comparing linguistic and acoustic predictors of canonical response components. **A**) We re-predict the canonical response components (obtained using the same decoder as in Figure 2) with a range of several feature spaces. **B**) Illustration of candidate feature spaces, segment shows 4 seconds of the speech stimulus. **C**) Test-set correlations for various feature spaces. Each data point is one outer fold of one participant. Thin horizontal lines show pooled means. Thick and thin vertical lines show 50% and 95% credible intervals of main effects of feature spaces from Bayesian model. See B for colour code. Large coloured dots on the x-axis denote feature space combinations, where data points are coloured according to the highest-level component. **D**) Statistical evaluation of main effects of feature spaces from Bayesian model. See B for colour code.

We were interested in two central questions: Are the neuronal dynamics we extract with our approach sufficiently rich to support a replication of recently reported correspondences of linguistic features with beta power modulations? Secondly, to what extent would we be able to explain such correspondences with acoustic predictors?

To answer our first question, we considered recent reports where power time courses in response to speech were modeled with syntactic features (Weissbart & Martin, 2024; Zioga et al., 2023).

In Zioga et al., (2023), these syntactic features describe sentence structures as directed links between individual words, referred to as dependencies. The authors characterised such dependencies with the opening, maintenance and resolution of arcs (Figure 3B). We could replicate the reported effect of an improved prediction performance when combining these three syntactic features with the original base model of Zioga et al. (2023) consisting of acoustic “edges”, word onsets and word frequency (Pearson correlation *ρ =* 0.2233) over the base model alone (*ρ =* 0.2076, Figure 3C). When comparing these models with a Bayesian approach, a fraction *f*_*h*_ *=* 0.9969 of the posterior samples for these two models supported this hypothesis (Figure 3D), translating into “decisive” evidence (Jeffreys, 1998).

In Weissbart & Martin (2024), the syntactic features were instead operationalised using constituency parsing, where words are grouped into hierarchical phrases whose relationships are then described. The authors extracted the depth in the parse tree as well as the amount of constituents that were finished (“Close”) at each word. Additionally, the authors extracted word-level surprise and entropy features from GPT-2, a language model (Figure 3B). In our canonical response components, the base model used in Weissbart & Martin (2024), consisting of a speech envelope and word frequency, achieved a performance of *ρ =* 0.2365. Adding only the constituency parsing features could only weakly improve on that (*ρ =* 0.2400, *f*_*h*_ *=* 0.7318, “barely worth mentioning”). Further adding GPT-2 features to this model however yielded a clearer increase in performance over the base model (*ρ =* 0.2424, *f*_*h*_ *=* 0.8415, “substantial”, Figure 3C,D). This mimics the original findings (Weissbart & Martin, 2024), where a combination of constituency parsing with GPT-2 features had shown the strongest improvements over the base model.

We next turned to our second question about the extent to which the performances of the high-level models could also be achieved with simpler acoustic features. Two models based on speech onsets (*ρ =* 0.2006) and gap onsets (*ρ =* 0.2057) were the worst models in our comparison. In line with the correlation between gap length and bursting duration described above (Figure 2G), a predictor indicating the presence of gaps however achieved a much higher performance (*ρ =* 0.2563). This outperformed all previously mentioned models with “decisive” (all *f*_*h*_ > .9932) or in the case of the model including constituency parsing and GPT-2 features “very strong” (*f*_*h*_ *=* 0.9846) evidence. It also outperformed other acoustic models such as the speech envelope (*ρ =* 0.2354, *f*_*h*_ *=* 0.9987, “decisive”) or the log-mel spectrogram in combination with its half-wave rectified temporal derivative (*ρ =* 0.2457, *f*_*h*_ *=* 0.9510, “strong”). Another way to create a gap-like predictor is to process the speech stimulus with a denoising algorithm (Défossez et al., 2020), removing any remaining sound energy during the gaps. In the case of our stimulus, which had been recorded in clean conditions, this mainly removed a stationary noise sound potentially coming from a fan close to the recording microphone. Consequently, the speech envelope computed on this denoised speech signal remained intact during speech, but emphasised the gaps more clearly (Figure 3B). Such a one-dimensional stimulus-computable feature can also be motivated from the observation that auditory cortex is invariant to noise sources (Khalighinejad et al., 2019; Mesgarani et al., 2014). It achieved a slightly better performance compared to the gap predictor (*ρ =* 0.2577, *f*_*h*_ *=* 0.5848, “barely worth mentioning”). Combining the denoised envelope with the dependency parsing predictors did not result in a performance gain (*ρ =* 0.2569), and the combination of the denoised envelope with constituency parsing features and GPT-2 features resulted in an improvement that was “barely worth mentioning” (*ρ =* 0.2593, *f*_*h*_ *=* 0.6020).

In summary, using cross-validated encoding models, we provided a more detailed characterisation of the canonical response component identified by our CCA analysis. Crucially, we could replicate a performance gain of syntactic dependency parsing features over a base model (Zioga et al., 2023) as well as a performance gain of a combination of “rule-based” and “statistical” features over another base model (Weissbart & Martin, 2024), demonstrating that our component was sensitive to previously reported high-level effects. However, we found simple acoustic features to be better predictors. Specifically, a one-dimensional predictor based on the sound energy of the denoised speech stimulus achieved the highest performance, which could not be substantially increased by combining it with the previously reported high-level predictors. The successful denoised speech envelope can be seen as a form of nonlinear compression, sharpening the contrast between sound energy during gaps and during the speech signal.

### Phase response of encoding models explains failure of generalisation testing to isochronous tones

Splitting neuroimaging datasets into train and test sets as done in (nested) cross-validation (Varoquaux et al., 2017) is a robust way to assess encoding or decoding models on “out-of-sample data” and common practise for many studies of neuronal responses to speech (Kim, 2022). However, it is rarely tested how well such models then generalise to other contexts, or “out-of-distribution”. Our previous finding of a simple acoustic feature predicting the same variance as high-level linguistic features suggests that the beta bursting reflects a domain-general auditory rather than a speech-specific process. In this case, our models should reproduce response patterns of beta power in other auditory experiments. Here, we turned to a simple experiment known from a landmark study on beta oscillatory responses. Fujioka et al. (2012) reported how beta power increased towards upcoming isochronous tones, before decreasing again after each tone. We extracted the grand average beta power responses to the four experimental conditions from the original paper (Figure 4A). We generated stimulus material as described in the corresponding methods section, fed these stimuli into our encoding models and correlated the model predictions with the original results. This allowed us to measure how well a given encoding model trained on audiobook listening data could account for results from a classic controlled experiment on predictable tone sequences at varying rates. Using our best feature space from previous analyses, the denoised envelope (see Figure 3), we found that this worked reasonably well for the random (mean performance *ρ =* 0.6246), slow (*ρ =* 0.7295) and medium conditions (*ρ =* 0.7619, Figure 4B, Figure 4J). For the fast condition however, generalisation performance was close to uniformly distributed, with a mean of *ρ =* 0.2951. Why would the models trained on speech not work in the fast condition of the experiment?

**Figure 4:**
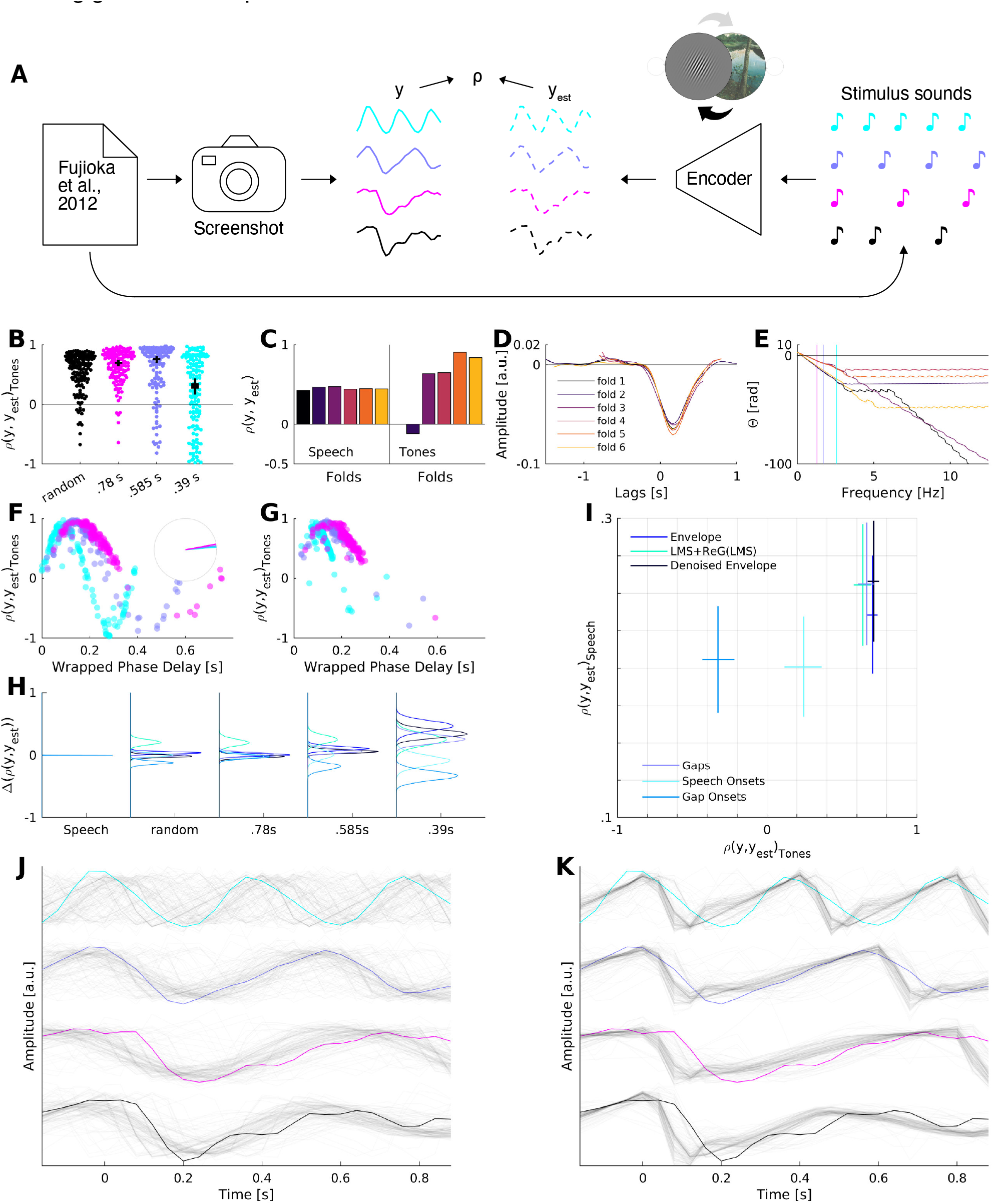
Encoding models trained with phase response regularisation on audiobook listening data can generalise to controlled rhythmic tones experiment. **A**) We extracted grand average beta power modulations in response to four experimental tone sequence conditions from Fujioka et al. (2012). Following the paper’s methods section, we generated corresponding stimulus waveforms and fed them into our encoding models. We then correlated extracted observations y and our model predictions y_est_. **B**) Generalisation performances in four conditions from Fujioka et al. (2012). Statistics from Bayesian model are overlaid in black, showing 50% (thick lines) and 95% (thin lines) credible intervals. **C**) Performances of encoding models of exemplary participant in audiobook listening and tones generalisation (fast condition). Here, performance on speech is highly similar across folds, but there is strong variability in generalisation performance. **D**) Encoding filters of all six cross-validation folds (exemplary participant) have highly similar shapes in time domain. **E**) The phase responses of the same filters as in D are however increasingly divergent at higher frequencies. Vertical lines indicate stimulus rates of isochronous tone sequence conditions. **F**) Wrapped phase delay (phase response measured in absolute time rather than as a phase angle) of encoding filters from all folds and all participants at frequencies corresponding to three isochronous conditions from Fujioka et al. (2012). E.g. the cyan dots are all intersections of phase responses in E (but from all participants) with the cyan vertical line. Inset shows angles corresponding to the best generalisation performance in all three conditions, which are highly similar. **G**) Same as F), but models are trained with an additional regularisation term constraining the phase response variance across frequencies. Most data points now cluster at similar and high-performing angles. **H**) Density estimates from Bayesian model comparing main effects of phase regularisation on performance of acoustic encoding models in speech (leftmost) and tones (right) data. Positive values reflect an improvement incurred by the phase regularisation. Speech performance is unaffected, while some models benefit most strongly in the fast condition. **I**) Comparing acoustic encoding models in 2D plane of performance in speech (y-axis) and tones (x-axis) data. Lines reflect 95% credible intervals of main effects of feature spaces from Bayesian model and intersect at the maximum-a-posteriori (MAP) of each distribution. **J**) Grand average beta power traces extracted from Fujioka et al. (2012) shown in solid colours corresponding to B, with predictions from denoised envelope model (all folds from all participants) shown as faint black lines. **K**) Same as J, but models are trained with phase regularisation term.

As our acoustic feature spaces are deterministic functions of the stimulus material, we turned to the temporal response functions (TRFs, i.e. the linear encoding models) as the only possible reason for this failure of generalisation. To our surprise, strongly varying generalisation performances across cross-validation folds (Figure 4C, right panel) were obtained with highly similar TRFs (Figure 4D). However, these TRFs are essentially finite impulse response (FIR) filters, which can not only be studied in the time domain, but also in terms of their phase response. This describes how the filter (i.e. the TRF) will delay the response at each frequency, and can be measured as a phase angle between input and output (de Cheveigné & Nelken, 2019). Interestingly, the same TRFs that looked virtually indistinguishable in the time domain exhibited nonlinear and non-constant phase responses which diverged markedly across folds especially for higher frequencies (Figure 4E) corresponding to faster conditions from the experiment. We attributed this to a lower signal-to-noise ratio of the neural response data at the faster frequencies, leaving little information to constrain the TRFs (Figure S1). Consequently, the TRFs’ phase responses were distributed close to uniformly at the frequency corresponding to the fastest condition (Figure 4F, x-axis, cyan points). High generalisation performance in all three isochronous conditions however was achieved at the same angle (Figure 4F, inset), while deviation from this angle resulted in a sinusoidally modulated performance (Figure 4F, y-axis). In other words, we found a strong relationship between the delay a TRF would incur to input at the frequency corresponding to the stimulus rate and the resulting generalisation performance.

We therefore devised a regularisation term to discourage the TRFs’ variance of their phase responses across frequencies (see Methods). Note that this did not constrain the TRFs’ phase responses to an absolute value. Training the TRFs for the denoised envelope feature with this additional regularisation clustered their phase responses in a single, favourable angle (Figure 4G). This did not measurably affect the performance of the encoding models on the speech data (Figure 4H, left panel). Crucially however, it drastically affected the generalisation performance especially in the fast condition for the envelope, the denoised envelope, the gaps and the combination of the log-mel spectrogram with its half-wave rectified temporal derivative (Figure 4H, right panel). In other conditions, the improvements were less strong, and other feature spaces worsened in performance (the speech onsets and the gap onsets). This now put us in the position to extend our model comparisons of our set of acoustic models to a 2D plane spanned by the performances of predicting *out-of-sample* responses to audiobook listening (Figure 4I, y-axis) as well as predicting *out-of-distribution* responses to rhythmic tone sequences (Figure 4I, x-axis, denoting the average across experimental conditions). Interestingly, the denoised envelope proved to not only be the best model to predict beta power modulations in response to speech, but also in response to rhythmic tones (*ρ =* 0.7176). While the gap model had been ambiguously close in performance on the speech comparison (see Figure 3C, D), evidence for a lower performance on the tones comparison (*ρ =* 0.6717) was “very strong” (*f*_*h*_ *=* 0.9857). The plain envelope achieved a performance on tones that was very close to that of the denoised envelope (*ρ =* 0.7052, *f*_*h*_ *=* 0.6640, “barely worth mentioning”), but had been beaten “decisively” by the denoised envelope in the speech comparison (see Figure 3C, D).

In sum, we here demonstrate the importance of testing models of brain responses not just on out-of-sample held-out data, but also in “out-of-distribution” contexts. We have shown that generalisation even to simple experiments is not trivial, but can be achieved by reducing hidden and unnecessary degrees of freedom of the encoding model.

### Testing models on passive speech listening data as well as controlled isochronous tones experiment improves model adjudication

Now equipped with a method to strongly improve the potential of encoding models to align with results known from the literature, we were next interested in applying the method to compare a broader range of feature spaces.

To define a set of feature spaces of interest, we firstly turned to two established deep self-supervised learning (SSL) speech processing architectures, wav2vec 2.0 (Figure 4A, Baevski et al., 2020) as well as contrastive predictive coding (CPC, Figure 4B, van den Oord et al., 2018). Both of these models yield versatile high-dimensional representations that are useful for a range of downstream tasks, and both models use a module of convolutional layers to downsample raw speech waveforms to slower, abstract latent states.

Wav2vec 2.0 (Figure 5A) then learns to contextualise these latent states with those extracted from both preceding and subsequent portions of the waveform using a stack of 25 transformer layers (see methods for details). We obtained activations from these transformer layers and used the 32 principal components explaining the most variance to predict beta power modulations.

**Figure 5:**
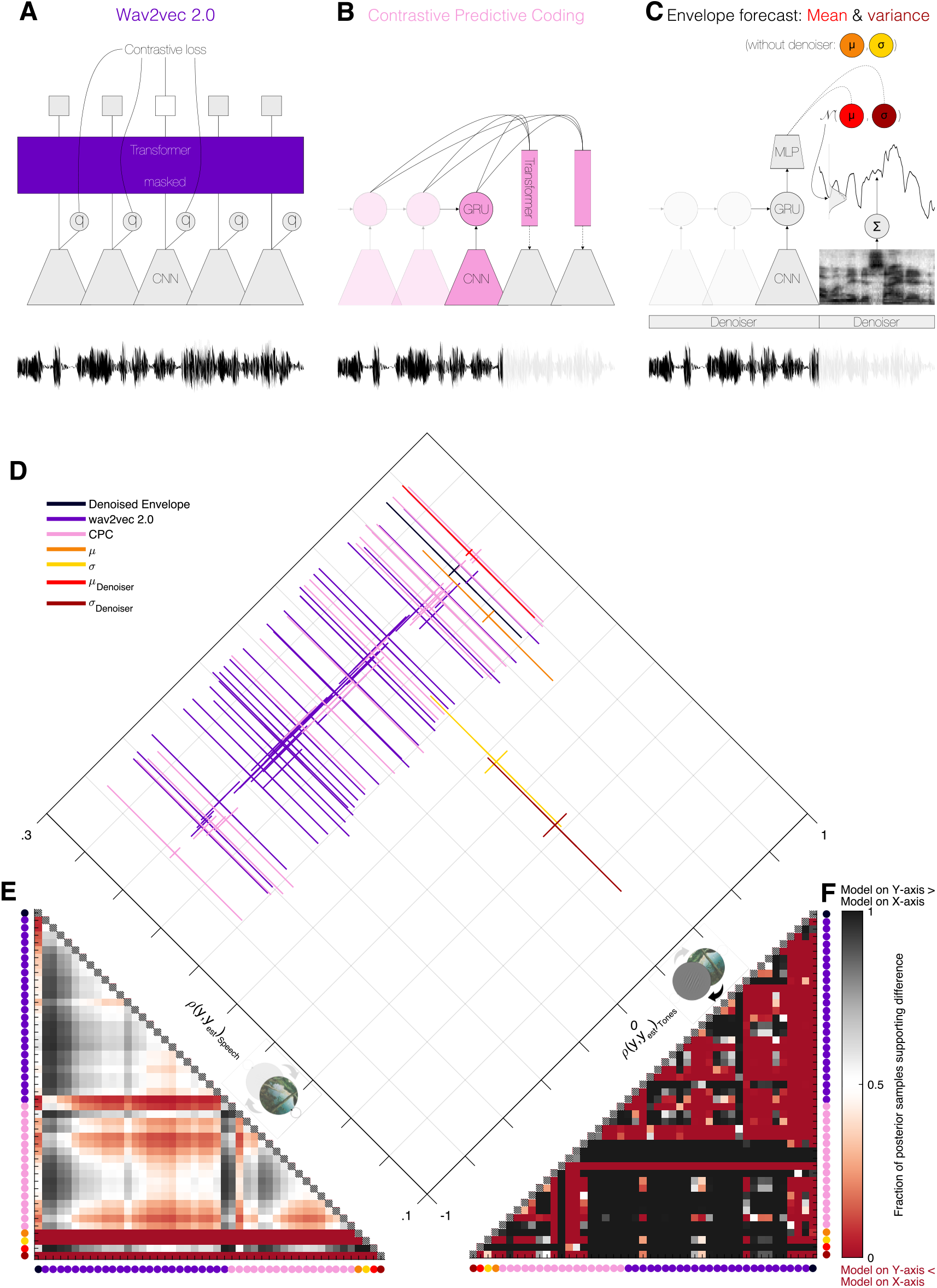
Adjudicating between models on naturalistic passive speech listening and controlled isochronous tones experiment. **A**) Wav2vec 2.0 architecture, adapted from Baevski et al., (2020). The network extracts abstract latent states from the waveform using a stack of 1D convolutions (convolutional neural network, CNN). It then contextualises these latent states with information from both preceding and subsequent latent states via a block of 25 transformer layers (purple). This is achieved with a masked contrastive loss during training, where the similarity between transformer outputs and the true quantised latent states (relative to other randomly selected latent states) is maximised. **B**) CPC architecture, adapted from van den Oord (2018). A gated recurrent unit (GRU) aggregates a sequence of CNN latent states and feeds into separate transformer layers that predict latent states extracted from future portions of the waveform. It is also trained with a contrastive loss (see methods). **C**) Custom probabilistic envelope forecasting model. The architecture is based on CPC, but simplified: The GRU feeds into a multilayer perceptron (MLP), which parameterises a Gaussian distribution of expected envelope values of the upcoming speech sample. We trained the model in two versions: Without a denoiser and with a denoiser preceding the CNN. **D**) Scatter plot showing test-set performances of encoding models trained on naturalistic passive speech listening data within-distribution (left axis) and out-of-distribution in tones experiment (right axis). The ideal model would appear in the top corner. Lines reflect 95% credible intervals of main effects of feature spaces from Bayesian model and intersect at the MAP of each distribution. 25 transformer layers of wav2vec as well as CNN, GRU and 15 transformer layers per forecast horizon from CPC are plotted individually, but in the same colour. **E**) Statistical evaluation of main effects of feature spaces from Bayesian model for speech comparison. See D for model colour code, see F for effect size colour code. Matrix rarely saturates, denoting mainly ambiguous evidence for model adjudication. **F**) Statistical evaluation of main effects of feature spaces from Bayesian model for tones comparison. See D for model colour code. Matrix mostly saturates, denoting mainly decisive evidence for model adjudication.

CPC on the other hand (Figure 5B) operates in a strictly causal fashion, aggregating a sequence of latent states with a recurrent module to then predict upcoming latent states with separate transformer modules for each of a range of forecast horizons (see methods for details). We used activations of the final convolutional layer and the recurrent layer as well as from each of the 15 transformer modules for forecast horizons predicting latent states up to .6 seconds ahead. We again reduced the dimensionality of these representations by keeping the 32 top principal components.

We added a third custom model (Figure 5C) which was based on the CPC architecture, but instead of predicting abstract latent states extracted by convolutional modules, the network simply learned to predict the upcoming sound energy. It did so by parameterising a Gaussian distribution, i.e. dynamically forecasting the upcoming envelope level (*μ*) and an associated uncertainty (*σ*). To match this model to our best acoustic feature, we also trained a version where a denoiser module preceded the convolutional encoders and the extraction of the upcoming envelope, yielding *μ*_*Denoiser*_ and *σ*_*Denoiser*_. Each of these four features were one-dimensional.

To compare these feature spaces, we considered the two-dimensional plane spanned by performance on predicting held-out test sets on the passive speech listening data as well as performance in generalising to the controlled isochronous tones experiment. We were interested in answering the following three questions: 1) Would the consideration of the tones experiment provide grounds to improve adjudication of the feature spaces beyond the speech listening data? 2) Would we be able to improve on the baseline provided by our best acoustic feature (see Figure 4I) with rich representations obtained from SSL models operating on the raw speech waveform? 3) Would our simplified and more interpretable custom model predicting only upcoming sound energy achieve a comparable performance?

Our first general observation was that most of the models considered here were close to the highest achieved performance on the speech evaluation (Figure 5D): A group of 44 models fell in the range from *ρ =* 0.2498 to *ρ =* 0.2576. This was not the case for the evaluation on the tones experiment, where performance ranged from strongly negative (*ρ =* −0.7673) to strongly positive values (*ρ =* 0.8089). Further, most model comparisons yielded ambiguous inferential statistics on the speech evaluation (Figure 5E, mostly non-saturated colours), whereas the majority of model comparisons in the tones evaluation indicated decisive evidence (Figure 5F, mostly saturated colours). The additional consideration of the tones experiment thus strongly improved our ability to adjudicate between models.

Our reference model in this comparison was the previously identified denoised envelope (see Figure 4I, speech comparison: *ρ =* 0.2568, tones comparison: *ρ =* 0.7103). One layer of the wav2vec 2.0 model achieved a higher performance on the tones comparison (layer 2, *ρ =* 0.7400, *f*_*h*_ *=* 0.9109, “strong”), but performed worse on the speech comparison (*ρ =* 0.2503, *f*_*h*_ *=* 0.9327, “strong”). Three transformer layers of the CPC model also achieved higher performances on the tones comparison than our best acoustic feature. These corresponded to forecast horizons of 5, 6 and 8 samples. 6 samples (i.e. .24 seconds) achieved the highest performance (*ρ =* 0.8089, *f*_*h*_ > 0.9901, “decisive”), and also came close to the denoised envelope on the speech comparison (*ρ =* 0.2552, *f*_*h*_ *=* 0.6415, “barely worth mentioning”). A layer of the causal, predictive SSL model (CPC) was thus an overall better model than our best acoustic feature in our comparison, while this was not the case for any layer of the post- and predictive SSL model (wav2vec 2.0).

The *μ*_*Denoiser*_ feature of our custom network predicting simply the upcoming sample of the denoised envelope came close to the best CPC performance on the tones comparison (*ρ =* 0.7969), translating into “substantial” evidence for the CPC model beating our custom model (*f*_*h*_ *=* 0.7824). However, this was reversed on the speech comparison, where the *μ*_*Denoiser*_ feature yielded the overall best performing model in the comparison (0.2576), which was “substantial” evidence for beating the best CPC forecast horizon (*f*_*h*_ *=* 0.7699), and “barely worth mentioning” in comparison to the denoised envelope (*f*_*h*_ *=* 0.5970). Neither the *μ* feature (i.e. without the denoiser) nor the variance features (*σ* and *σ*_*Denoiser*_) made it into the set of competitive models. Aggregated across both comparisons, our simple custom network was thus competitive with the best SSL feature space.

Taken together, we found that while many candidate models performed similarly on the speech evaluation, the additional consideration of the simple isochronous tones experiment helped to tell the models apart. We thus learned that our acoustic baseline (the denoised envelope) could be beaten with high-dimensional activations from a general-purpose SSL neural network trained to predict abstract latent states extracted from the upcoming speech signal (CPC). Crucially, a one-dimensional feature from a highly simplified network predicting simply the upcoming loudness was competitive with the CPC model. We thus showed how a computationally explicit approach of bridging between findings from uncontrolled naturalistic datasets to controlled experimental settings known from the literature allowed us to better adjudicate between our models. This marks an important step towards a more integrative approach to neuroscience (Schrimpf et al., 2020).

### Network advantage over acoustic predictors is rooted in slow decay

Lastly, we were interested in better understanding how our custom network model achieved its advantage over our best acoustic model. Was this due to yet another trivial form of generic nonlinearity applied to sound energy (Daube et al., 2019)? Or was this instead genuinely attributable to its autoregressive training objective (Antonello & Huth, 2024)? To do so, we aimed at localising specific moments in time that contributed to the higher performance of our network feature over the acoustic feature. We did this by subtracting the absolute values of the residuals of both predictions, such that positive values would correspond to smaller residuals of the network model and negative values to smaller residuals of the acoustic model.

We first focused on the speech comparison. As expected from the small differences in performance between the network feature and the acoustic feature (see Figure 5), their predictions were highly correlated (average across folds and participants *ρ =* 0.9875, Figure 6A) and thus scattered close to the diagonal. The differences of absolute residuals (colour coding) did not yield a clear pattern on these predictions.

**Figure 6:**
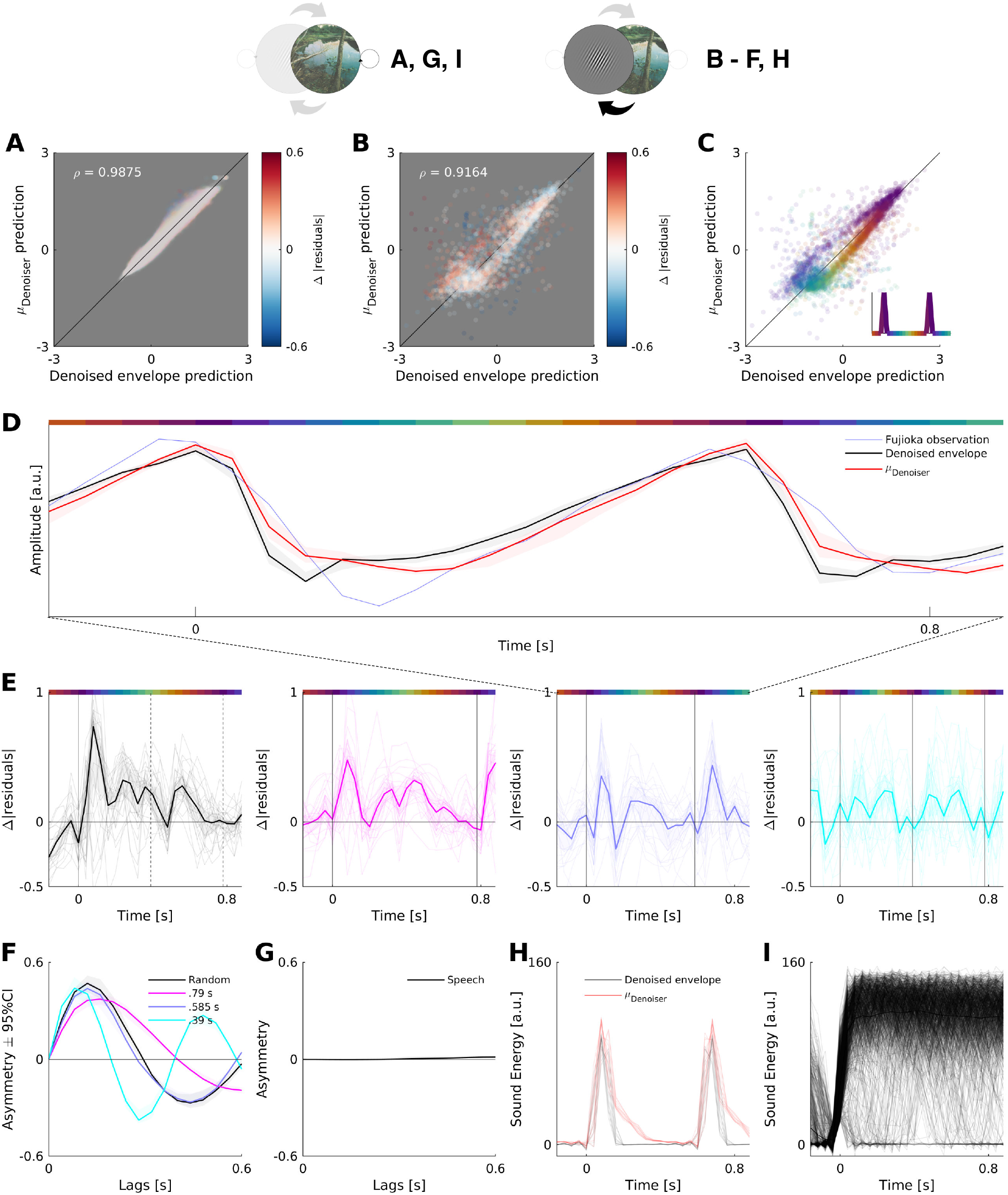
Localising the advantage of the network predictions over the acoustic predictions. **A**) Scatterplot of predicted neural responses to speech from two predictors. Colour denotes difference of absolute residuals (symmetric logarithmic scale), where red regions highlight lower residuals (better predictions) from network model. Data from all participants and all folds overlaid. Top left reports correlation of predictions, averaged across folds and participants. Predictions are highly correlated, residuals give no clear pattern. **B**) Scatterplot of predicted neural responses to tones from two predictors. Colour denotes difference of absolute residuals as in A. Data from all participants and all folds are overlaid. We show data from the medium condition (.585s), see Figure S2 for other conditions. Top left reports correlation of predictions averaged across folds, participants and all conditions. Compared to speech, the tones yield less correlated predictions with smaller residuals for the network feature in most data points. **C**) Same as in B, but colour now denotes time relative to tone onset (see inset for legend), showing stimulus structure in the scatter plot. **D**) Observed and modeled beta power dynamics surrounding tone onset. See C for colour code of line at the top. **E**) Differences in absolute residuals for all four experimental conditions. Solid lines show averages across folds and participants, faint lines show individual folds and participants. Vertical lines denote tone onsets. Positive values denote lower residuals from network model. Insets show average differences of raw predictions (not residuals) mapped onto observed neural responses from Fujioka et al. (2012). Network always predicts higher values during beta power decay (purple) and lower values in rise to next peak (green), temporally aligned with lower residuals for the network. **F**) Asymmetries of cross-correlations (difference of cross-correlation functions over positive minus negative lags) between predictions from network feature and acoustic feature. Lines denote averages across folds and participants, shaded regions denote 95% confidence interval. Features are positively correlated at zero lag, positive values of the asymmetry at short non-zero lags thus denote a delay of the network features relative to the acoustic features, the sign then flips after zero-crossings of the cross-correlation functions. Strong asymmetries are visible in all four conditions. **G**) Same as in E, but for speech data. No clear asymmetries are visible. **H**) Raw acoustic feature and network feature in response to tones (medium condition, see Figure S2 for other conditions). Each line is a single trial. Network output decays slower than acoustics. **I**) Segments of denoised envelope following gaps in audiobook. Sound energy is mostly sustained following gaps, and rarely decays quickly.

This was different for the tones comparison, where we had observed stronger differences in performance (see Figure 5). Here, the predictions based on the two features deviated from a simple linear relationship (average across folds, participants and conditions *ρ =* 0.9164) and instead scattered as ellipses around the diagonals (Figure 6B, see Figure S2 for data from all conditions). These ellipses were structured according to time relative to tone onsets (Figure 6C, see Figure S2 for data from all conditions).

When considering the same data as event-related traces, we could see how the network model was closer to the observation than the acoustic feature across most of the time surrounding the tone onset (Figure 6D, medium condition, see Figure S2 for all conditions). Two moments particularly drove this advantage, yielding two peaks of the differences of absolute residuals in favour of the network model in all 4 conditions (Figure 6E): Shortly after the onset of a tone, i.e. during the decrease from the peak beta power, and during the rise towards the next peak. The more accurate network feature was predicting higher values than the acoustic feature during the decrease of power, and lower values than the acoustic feature during the following rise. In other words, the network feature predicted a trajectory that was delayed relative to the acoustic feature, which was discernible in clear asymmetries of the cross-correlation functions of network and acoustic predictions (Figure 6F). These asymmetries were not observable in the predictions of the responses to speech (Figure 6G).

To explain this, we compared the raw network outputs in response to the sound energy in the controlled tones experiment (figure 6H, see Figure S2 for data from all conditions). We found the main difference between the two to be in how they behaved following the tone: While the sound energy decayed quickly, the network had learned to predict a slower decay. We attribute such a “slow decay prior” to the data the network was trained on, which consisted of audiobooks. In such speech data, fast decays of sound energy following an onset are rare – onsets are mostly followed by sustained speech activity (Figure 6I). The interaction of such slowly decaying features with the corresponding TRF encoding models thus produced trajectories that stuck to the high peak values for longer, decayed at a later time and then started to ramp up towards the next onset later. This achieved a higher performance in predicting the experimental data observed in Fujioka et al. (2012).

Taken together, our results demonstrate again how considering the controlled tones experiment improved model adjudication by decorrelating predictions from different models. In this way, we show how the advantage of our forecasting network over our acoustic features is rooted in the internalisation of a “slow decay” statistic learned from a large audiobook dataset. The advantage of our forecasting network is thus not simply a consequence of e.g. a trivial compressive nonlinearity applied to acoustic energy, but instead resulting from correctly predicting a more sluggish response to fast-decaying transient sounds.

## Discussion

Are old experimental designs based on simple artificial stimuli still relevant in the modern age of naturalistic neuroscience, or are they merely historic artefacts? We show here that existing literature is in fact a treasure trove of diagnostic benchmarks that should be exploited to more severely test modern encoding models beyond naturalistic stimuli. We demonstrate this on the example of beta power dynamics in response to audiobook listening. While we can replicate reported links of this understudied response component to high-level linguistic parsing, we find that a parsimonious acoustic model explains the same variance in our MEG responses. This suggests that beta responses may reflect a domain-general auditory process rather than one specific to language. Models should thus generalise beyond speech to other acoustic stimuli.

We find that such backwards compatibility to a classic experiment based on rhythmic tone sequences is possible, but only with a novel regularisation of the phase response of speech-trained encoding models. This then allows us to compare a wider range of deep stimulus-computable models on both passive audiobook listening data and the controlled tones experiment, improving our capacity to adjudicate between models: They perform similarly on predicting held-out responses to speech but differ greatly in their predictivity of responses to pure tones. In this way, we find that activations from an autoregressive self-supervised deep neural network model beat our strongest acoustic baseline. Further, we find that we can achieve a comparable performance with a simpler custom network trained to simply predict the upcoming sound energy. Finally, we show how this custom network derives its tone-specific advantage over acoustic models from a slow-decay prior learned from speech statistics. These results demonstrate how the field should move beyond model comparisons limited to a single stimulus class, and instead embrace an integrative and computationally explicit approach for model development and testing.

### High-level versus acoustic predictors of beta power responses

In this study, we successfully replicated previous reports of performance gains of high-level linguistic features over acoustic features in our response component. However, similarly as in previous work (Daube et al., 2019), we then found a simpler feature – the speech envelope computed on the denoised speech signal – to capture the same response variance: The test-set prediction performance of the denoised envelope could not be substantially improved by combining it with the high-level features. Of note, our best predictor was thus competitive despite consisting of just a single dimension, beating concatenations of six unidimensional features. What does this imply for the interpretation of power responses during speech listening?

In the same way that the original reports do not conclusively prove that responses under study are in fact representing the high-level processes (Guest & Martin, 2023), our finding of features that predict the same variance is not conclusive proof of the contrary. While our stimulus-computable features have the allures of (i) low dimensionality, (ii) conceptual simplicity in comparison to syntactic or predictive processing, (iii) mechanistic explicitness and plausibility for processes implemented in the auditory pathway as well as (iv) connections to previous findings of noise-invariant speech representations in auditory cortices (Khalighinejad et al., 2019; Mesgarani et al., 2014), our results remain correlational. We do not dispute that humans are able to perform the cognitive operation of tracking syntactic structures, or that humans can be more or less surprised by an incoming word given its context, as demonstrable with behavioural dependent metrics (Krakauer et al., 2017; Marslen-Wilson & Tyler, 1980; Miller & Chomsky, 1963; Miller & Isard, 1964; Niv, 2021). We further consider linguistic ideas about neuronal speech processing an invaluable source of hypotheses about the functional significance of observable responses (Gwilliams et al., 2025; Poeppel & Embick, 2005) that inspires further research and yields benchmarks for stimulus-computable process models.

We however put into question whether the limited view we get of the listening brain via (non-invasive) neuroimaging necessarily provides access to the neuronal underpinnings of these computations. To move forward, controlled experiments (Ding et al., 2016; Schyns et al., 2022) that move beyond “naturalistic” audiobook listening by equalising all relevant acoustic stimulus properties are thus needed.

Ultimately, we hope for a fruitful exchange between approaches, in which some researchers will continue to describe situations where high-level features beat acoustic baselines and therefore challenge bottom-up models, whereas other researchers search for stronger acoustic baseline features, which are ultimately also needed to deconfound the required controlled experiments.

### Predictive models win because of a slow-decay prior

Beta band activity has previously been portrayed as being related to temporal forecasting (Arnal et al., 2015; de Lange et al., 2013; Grabenhorst et al., 2025; Sedley et al., 2016). We therefore considered autoregressive models as candidates to potentially beat the performance of our best acoustic model in the joint evaluation on the speech dataset and the tones experiment. We indeed found that individual layers of CPC, a DNN that forecasts future latent states extracted from the speech waveform, outperformed our best acoustic model on the tones evaluation. Individual dimensions of the representations extracted from the CPC model are however difficult to interpret, leaving room for alternative explanations based on e.g. the effective dimensionality of model activations (Elmoznino & Bonner, 2024). We thus developed a custom network simply forecasting the upcoming sound energy (and an associated uncertainty, Obleser, 2025) from two seconds of the raw waveform. In our comparisons, the prediction of the expected value of the upcoming sound energy was on par with the CPC model, providing a much simpler and uni-dimensional narrative: The beta bursts appear to be captured well by a forecasting process that attempts to anticipate loudness variations.

This however leaves open the question: Are autoregressive optimisation objectives just “one way out of many to discover useful […] features” (Antonello & Huth, 2024) for predicting beta bursts? A potential counter-argument is that in our joint evaluation, activations extracted from any of the layers of another self-supervised but non-autoregressive model (wav2vec 2.0) failed to outperform our best acoustic model.

Further, we revealed the concrete property that allowed our predictive models to beat acoustic models in the tones comparison. By studying the model predictions at a finer scale, we showed that the advantage of the mean-prediction network model over the best acoustic model was rooted in an internalisation of a “slow-decay” prior the model had likely learned from audiobook speech data in order to minimise the autoregressive cost function. Following onsets, speech signals usually do not decay instantaneously, and so an effective way to optimise the loss function is to default to a slow decay. When confronted with the fast-decaying artificial tones, following this prior is not adaptive, but helpful to predict beta power dynamics in response to the same tones. We therefore put forward that the human auditory system might be similarly overfit to sluggish speech dynamics.

### Comparing biological systems and models on naturalistic or artificial stimuli

A central result of our study is the low discriminability between many high performing models when using only the uncontrolled speech data. Crucially, the additional testing of backwards compatibility to a controlled experiment from the literature (Fujioka et al., 2012) revealed strong performance differences. This links our results to an increasingly ubiquitous problem in cognitive computational neuroscience, where many different computational models achieve the same level of performance when compared with respect to uncontrolled naturalistic stimuli (Conwell et al., 2024; Golan et al., 2020; Putnam, 1967).

If the goal of the neuroconnectionist research programme (Doerig et al., 2023) is to identify the best model of human brain function, this common practise of exclusively relying on uncontrolled stimulus material such as scene images or audiobooks will fail (Bowers et al., 2022; Schyns et al., 2022). In order to better adjudicate between different models, the additional consideration of artificial stimuli as used in existing literature (Bowers et al., 2022) or as obtained from clever model-specific stimulus optimisation procedures (Feather et al., 2023; Golan et al., 2020) is required. Such artificial stimulus material can test hypotheses about generalisation gradients along specific stimulus parameters (Daube et al., 2021; Leske et al., 2025; Rust & Movshon, 2005; Schyns et al., 2023), enhancing interpretability and providing inspiration for model improvements.

We however stress that this does not imply that comparisons on naturalistic stimuli are uninformative: Sensory neurons are assumed to be tuned to naturalistic statistics, and so their responses typically become more vigorous (Rieke et al., 1995) and can change nonlinearly when presented with naturalistic as opposed to artificial stimuli (David et al., 2004; Theunissen et al., 2000).

As in the study at hand, tests are thus ideally performed on both exploratory naturalistic and confirmatory artificial domains (Oganian & Chang, 2019; Tukey, 1980). This incentivises the development of computationally explicit models that integrate across seemingly disparate subdomains of sensory neuroscience (Schrimpf et al., 2020). As we show here, importing experimental results from the literature to naturalistic projects can be strikingly simple by directly obtaining data from figures of published papers. In doing so, we establish a direct link between beta bursting observed during simple tone trains and those observed during speech listening. This direct link is not at all trivial, as successful generalisation is only achieved by incorporating a phase-constancy prior on the encoding models. By making this link computationally explicit, we make a strong case for the underpinning mechanism of the beta bursting to be primarily related to domain-general acoustic stimulus dynamics.

### Constraining phase responses of encoding models

While it is common practice to examine phase responses of spectral filters (de Cheveigné & Nelken, 2019) and also widely appreciated that linear encoding models based on the formalism of temporal response functions are in essence finite impulse response filters (Crosse et al., 2016), it is to our knowledge uncommon to consider the phase response of TRF encoding models. Here, we find that the phase responses of our naively trained linear encoding models exhibit strong and variable nonlinearities, even for instances that appear highly similar in the time domain. Interestingly, we then find that while constraining this flexibility does not harm out-of-sample (but within-distribution) prediction performance, it greatly boosts the performance of generalising to out-of-distribution data. This adds to mounting accounts that describe linear reweighting of feature spaces as an overly flexible tool to assess their brain-likeness (Conwell et al., 2024; Lampinen et al., 2025; Schaeffer et al., 2024).

The generalisation is achieved without controlling the absolute phase angle between the stimulus and the response, but simply by encouraging a constant phase angle across frequencies. The constrained phase response thus essentially affects how the encoding model deals with stimulus rates not encountered in the audiobook stimulus. We assume that successfully generalising encoding models could similarly be obtained by training them on MEG data recorded while listening to speech stimuli covering a range from slow to fast rates (Ahissar et al., 2001).

Our results suggest that the beta bursts occur at a constant low-frequency stimulus phase angle across speech rates. Since transient stimulus events occurring at a matching rate have in turn been shown to be in a relationship with neural delta phase (Chalas et al., 2023), this fits observations that describe beta bursting as nested within neural delta phase (Keitel et al., 2018; Weissbart & Martin, 2024). It also fits the finding that neural delta phase is in a phase relationship with the stimulus that is constant across frequencies (Doelling et al., 2019). However, the faster neural theta phase has been shown to be in a non-constant linear phase relationship with the stimulus (Zou et al., 2021). It is thus an open question how beta bursting dynamics will behave for faster stimulus rates than what was tested in acoustic experiments (Fujioka et al., 2012; Pesnot Lerousseau et al., 2021), or in paradigms using time-compressed speech (Ahissar et al., 2001; Giroud et al., 2024): Beta bursting might stop demarcating stimulus onsets entirely, or its phase relationship to the stimulus might change.

An important neuroscientific question concerns what the functional significance of such constant phase responses could be. A property of neuronal populations that changes as a function of the phase of low-frequency oscillations is neuronal excitability (Buzsáki & Draguhn, 2004; Haegens et al., 2011). Neuronal excitability could create the necessary conditions for beta bursting to happen (Henry & Herrmann, 2014), such that metabolically costly bursting would be restricted to brief moments in time, potentially to sample relevant parts of the input at higher fidelity (Schroeder & Lakatos, 2009). Correspondingly, beta bursting nested into specific low-frequency phase angles has been shown to be in a positive relationship with the accuracy of perceptual judgements of auditory stimuli (Arnal et al., 2015; Herrmann et al., 2016). More generally, a restriction of beta bursting to relevant and attended stimulus details (Saleh et al., 2010) would also be in line with a recent EEG study where a deep neural network was employed to decode spatial attention to speech streams coming from the left or the right (Vandecappelle et al., 2021): Interestingly, the authors found that removing beta band activity with a corresponding notch filter resulted in the strongest decrement of predictive performance. A relationship of beta dynamics and attention further fits with a recent suggestion whereby the beta band sits in a frequency range ideal for alternating between bottom-up and top-down processing streams, achieving a balance optimal for the identification of syllables (Hovsepyan et al., 2023). We did not attempt to model the phase of beta signals in this study, but suggest that the power dynamics we describe in the study at hand could manage the deployment of such more detailed beta-phase-specific processes.

### CCA as a sensitive and computationally efficient joint encoding and decoding model

The lynchpin of our analysis is CCA, essentially implementing a joint linear decoding and encoding model (de Cheveigné et al., 2018; Dmochowski et al., 2018; Wang et al., 2020). This exploits the covariance structures of both a multivariate response space, here consisting of power time courses from multiple spectral bands at multiple MEG sensors, and a multivariate stimulus space, here consisting of lagged versions of the speech envelope. It therefore improves the sensitivity to find correspondences between neuronal responses and stimulus features beyond isolated linear encoding or decoding models. This is comparable to a previous approach (Daube et al., 2019) in which we obtained “decoding” weights as spatial filters of bilateral dipoles. In that case, we subjected the location and orientation of these dipoles to a black-box hyperparameter optimisation routine that we also used to optimise other hyperparameters such as the amount of regularisation of an encoding model. Our earlier approach can thus be seen as making use of the biophysical priors of the spatial filtering algorithm, but in turn assumes a locality of the effect of interest in MEG source space (Gwilliams & King, 2020). CCA, as used in the study at hand, relaxes this assumption and linearly combines sensors in a more data-dependent way.

For one-dimensional stimulus features, this is in principle also implementable as a pure decoding analysis. However, such an implementation would require computing a covariance matrix (and a singular value decomposition thereof) over combinations of MEG sensors, spectral bands and lags, resulting in a severe computational bottleneck.

As each sensor and frequency could in this rather theoretical case have individual weight profiles over lags within a single component, it is possible that such an approach would be able to pull out more or richer response components than the single component we found. However, even in this single projection we could replicate most of the previously reported high-level effects. Crucially, this single component was sufficient to generalise our findings to a controlled experiment from the literature, a central goal of our study.

We thus endorse CCA as a pragmatic and powerful alternative to isolated encoding or decoding models, and point out that further more detailed characterisations of power responses during speech listening should be explored in future research.

### Conclusion

We here proposed a canonical temporal forecasting model of beta power dynamics during listening. We first designed a robust method to extract beta power dynamics from sensor-level MEG data and replicated previous reports asserting beta power to be related to high-level linguistic processing with the extracted response component. We then found a simple, sound-energy-based feature that predicts the same response variance and established a computationally explicit link that makes our model backwards compatible from naturalistic speech to rhythmic tone stimuli known from the literature. We next assessed a range of DNNs in our two-dimensional performance space of predicting beta dynamics during speech and tone listening. By virtue of our enriched model comparison, we found that some DNNs indeed beat our simple acoustic feature. Specifically, we saw that the competitive performance of a self-supervised DNN that forecasts complex future latent states could also be achieved with a simple DNN that merely forecasts the upcoming sound energy. This network internalises a slow-decay prior from speech statistics, which allows it to outperform acoustic models. Taken together, our results highlight that the field should move on from the common practice of evaluating models within-distribution, and instead turn to testing on a broader set of out-of-distribution test cases.

## Acknowledgements

We thank the labs of Benjamin Morillon and David Poeppel for helpful discussions and Sander van Bree for helpful comments on the manuscript. The authors received no specific funding for this work.

## Author contributions

C.D. conceived of and designed the study. C.D. collected and analysed the data. C.D. and R.A.A.I. contributed analytic tools. C.D. wrote the initial draft. C.D., R.A.A.I., and J.G. edited the manuscript. R.A.A.I. supervised the project.

## Data availability

We will release the MEG dataset in an open repository upon publication. The librispeech dataset is openly available at https://www.openslr.org/12.

## Code availability

We have uploaded the code to github repository which will be made publicly accessible upon publication.

## Methods

### 1.1 MEG recordings

This study used a dataset that had previously been recorded (Daube et al., 2019) and analysed (Chalas et al., 2022, 2023; Daube et al., 2022). In the experiment, 24 native speakers of English (12 female, mean age 24.0 years, age range 18 - 35 years) had provided informed written consent and received a compensation of £9 per hour. The experiment had been approved by the Ethics Committee of the College of Science and Engineering (application number: 300170024).

The speech stimulus used in the experiment had a total duration of 55 minutes (“The Curious Case of Benjamin Button”, public domain recording by Don W. Jenkins, librivox.org) and was presented in 6 blocks of equal duration, where the last ten seconds of the first five blocks were presented at the start of the following blocks as a lead-in to maintain narrative continuity for the participants. We used MEG compatible Etymotic ER-30 insert earphones to deliver the sound, while participants sat in a magnetically shielded room where their brain activity was recorded with a 248 magnetometer MEG system (MAGNES 3600 WH, 4D Neuroimaging) at a sampling rate of 1017.25 Hz (first 10 participants) and 2034.51 Hz (last 14 participants). Each participant’s head shape was digitised, and five head position measurement coils were attached to bilateral pre-auricular points as well as three positions spread across the forehead. We measured the head position with respect to the sensor array before and after each recording block and repeated a block if the movement exceeded 5 mm. Participants demonstrated that they had attentively listened to the story by answering 18 multiple choice questions with 3 options each, with an average accuracy of .95 (standard deviation [SD] .05, range from .78 to 1.00).

### 1.2 MEG preprocessing

We used the same preprocessing pipeline that had been applied previously (Daube et al., 2019), which had made use of the fieldTrip toolbox (Oostenveld et al., 2011). Data was epoched using the start of the blocks (including the lead-in) as a trigger. We subtracted the projection of the raw data onto an orthogonal basis of the reference channel from the raw data to attenuate noise. Artifactual channels (3.07 channels per block on average, SD 3.64) were manually replaced with spherical spline interpolations of surrounding channels, and squid jumps were replaced with DC patches. We filtered the data with a 4th-order butterworth forward-reverse zero-phase high-pass filter with a cutoff-frequency of .5 Hz and subsequently discarded the lead-in parts from blocks 1 to 6. We downsampled the data to 125 Hz to run an independent component analysis using the *runica* algorithm. We manually identified artifactual components suggestive of heart or eye activity (6.70 components per block on average, SD 5.01), unmixed the data at the original sampling rate and backprojected it using mixing matrices where artifactual components were removed. We next downsampled the cleaned data to 100 Hz and processed it with a bank of 20 3rd order butterworth forward-reverse zero-phase bandpass filters, with lower cut-off frequencies starting at 1 Hz and increasing in steps of 2 Hz to 39 Hz, and upper cut-off frequencies starting at 4 Hz and increasing in steps of 2 Hz to 42 Hz. We obtained the absolute values of the hilbert transform of the filtered time series at each sensor and downsampled this to 25 Hz, resulting in a (time x [sensors * frequencies]) 82500 x [248 * 20] matrix per participant.

## 2 Feature spaces extracted from speech stimulus

We extracted a range of acoustic, linguistic and deep neural network based features from the speech stimulus.

### 2.1 Acoustic features

The basis for our acoustic features was a 31-channel log-mel spectrogram (LMS) extracted from the waveform of the audiobook, with centre frequencies ranging from 124.1 Hz to 7284.1 Hz. We used the implementation provided in the GBFB Matlab toolbox (Schädler et al., 2012). To obtain a 1-dimensional representation of time-varying sound energy (“speech envelope”), we summed the LMS across bands. For an operationalisation of acoustic onsets, we combined the LMS with its half-wave rectified temporal derivative. To replicate previous findings (Zioga et al., 2023), we also obtained a 1-dimensional time course of “edges”, defined as the 97.5th percentile of the temporal derivative of the speech envelope.

To test models focusing on the presence or absence of speech sounds only, we defined gaps as brief pauses in the audiobook. We did this based on the observation that the speech envelope is bimodally distributed, with one peak reflecting the speech energy during voice activity, and with another peak at low energy reflecting background noise during pauses, e.g. between sentences. Although the audiobook was recorded in “clean” conditions, this background noise is not completely silent, since it includes e.g. static fan noise of the computer used to record the audiobook. We defined gaps as moments where the envelope dropped below the trough between the two peaks in the distribution of the envelope. From this, we defined a “gaps” time series with ones during gaps and zeros everywhere else, a “speech onsets” time course with a single unit impulse at the end of gaps and a “gap onsets” time course with a single unit impulse at the start of gaps.

Finally, we also applied a speech denoising algorithm (Défossez et al., 2020) to the audiobook and recomputed feature spaces derived from the LMS on this denoised version. We found this denoiser to improve in performance when applied iteratively, where we added Gaussian noise with an amplitude of .003 to the waveform prior to running the denoising algorithm, which we repeated six times.

### 2.2 Linguistic features

To replicate previous findings (Zioga et al., 2023), we extracted a range of linguistic features from our stimulus. To do so, we re-used an annotation of the acoustic stimulus that had previously been created (Daube et al., 2019). Here, the Penn Phonetics Lab Forced Aligner (Yuan & Liberman, 2008) had been used to temporally align the text material to the stimulus waveforms, providing us with word identities and onset times. These had been manually corrected using Praat (Boersma, 2001). We created a word onset predictor directly from the onset times by populating a vector of zeros with unit impulses at samples that coincided with an onset of a word.

#### 2.2.1. Word Frequency

We used the wordfreq python library (Speer et al., 2016) to obtain word frequencies of words appearing in the story. We replaced small frequencies with a minimum value of 1e-9 and computed the negative logarithm of the result. Finally, we replaced the unit impulses of the word onset predictor with these values.

#### 2.2.2. Dependency parsing

We followed a previous report (Zioga et al., 2023) to extract features describing syntactic operations. We used the stanza python package (Qi et al., 2020) to parse the text of our speech stimulus, creating dependency graphs for each sentence. Then, we looped through the 593 sentences of the story. Within each sentence, we looped through each word. Punctuation symbols were ignored. If stanza produced a multi-word token (e.g. “*don’t”* becoming “*do”* and *“n’t”*), we inspected each component separately.

For each word following a given reference word, we added it to the number of nodes opened at the reference if the word was the head of the reference or vice versa.

If a word followed a reference word and had a head that appeared prior to the reference, or if a word preceded a reference word and had a head that followed the reference word, we increased the counter of nodes that remained open at the reference word. Stated differently, this tracked the number of nodes whose endpoints were on opposite sides of the current reference word.

Lastly, if a word preceded a reference word and either the reference was the head of the word or vice versa, this counted as a resolved node.

We replaced the unit impulses of the word onset predictor with the counts derived for opened nodes, nodes remaining open and resolved nodes, respectively.

#### 2.2.3 Constituency parsing

A further previous report (Weissbart & Martin, 2024) used Constituency parsing to describe syntactic operations. We again used the stanza package (Qi et al., 2020) with its implementation of a shift-reduce parser (J. Liu & Zhang, 2017) to obtain phrase-structure trees for the sentences of our speech stimulus. Following Weissbart&Martin (2024), we measured the stack “Depth” at each word in a left-to-right traversal. In other words, this tracks how deep into nested phrases the parser is at each successive word, and thus returns higher values when one enters a new constituent and lower values when one exits back up the tree. We also measured the amount of constituents that were finished at a given word, producing a “Close” predictor. Stanza parses each sentence starting with a root node and a sentence node. We discarded the root node and reassigned closings of the sentence node to the last lexical word in a sentence (instead of the final full stop punctuation mark).

#### 2.2.4 Language model features

Finally, following Weissbart&Martin (2024), we also extracted predictors describing the surprisal and entropy associated with the prediction of an upcoming word. To do so, we used a pretrained GPT-2-XL language model (Radford et al., 2019) as available from huggingface (Wolf et al., 2020). This model consists of 48 transformer blocks that operate on word embeddings of a dimensionality of 1600 and a maximal context window *l* of 1024 byte-pair encoding tokens. We propagated the text of the story through this model using a rolling context window of the maximal length afforded by the model. We obtained the logits of each final discrete token prediction and computed the log-softmax of these logits.

We then obtained the negative log-probability of token *t* assuming the actually observed identity *i* conditional on the prediction made based on the context comprising tokens *t* − 1, *t* − 2, *t* − 3, … *t* − *l* and stored this at the position of token *t*. We then computed the joint negative log-probabilities *S*(*w*) of individual words by summing the token-wise negative log-probabilities for all tokens *t*_1_, *t*_2_, *t*_3_, …, *t*_n_ comprising a given word *w*:

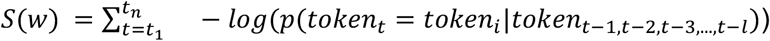

Assuming a positive constant delay (across words) from word onset of word *w* until the end of the hypothesised neurocognitive surprisal computation, we replaced the unit impulses at word onsets with the joint log-probabilities of that word.

We also computed the entropy *H* of the prediction of token *t* based on the context comprising tokens *t* − 1, *t* − 2, … *t* − (*l* − 1) by obtaining the negative sum (over the size D of the dictionary of tokens) of the product of probabilities and log-probabilities. For each word *w*, we kept the entropy after observing the first token *t*_)_ of that word:

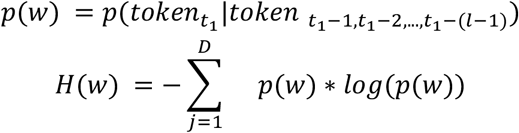

This again assumes a constant delay from the word onset of word *w* until the peak of the hypothesised neurocognitive state of uncertainty after having observed word *w*.

### 2.3 DNN-based speech features

#### 2.3.1 Self-supervised learning

Over the recent years, progress in self-supervised learning has yielded a range of models that can transform raw speech waveforms into rich abstract representations useful for downstream tasks. These have also been shown to be useful for predicting brain responses (Anderson et al., 2024; Millet et al., 2022; Vaidya et al., 2022). We therefore considered two models with respect to their capacity to predict our response component.

##### 2.3.1.1 Contrastive Predictive Coding

Firstly, we considered Contrastive Predictive Coding (CPC, van den Oord et al., 2018). CPC learns representations by predicting future latent states computed from the waveform by a convolutional encoder. It uses a contrastive loss encouraging the model to maximise the similarity between predicted latent states and true upcoming latent states and minimise the similarity between predicted latent states and randomly selected latent states. We modified an implementation of this model provided by facebook research (Riviere et al., 2020). The original implementation uses an encoder consisting of 5 layers of 1D convolutions, to which we added 2 more layers to achieve a downsampling factor of 640, such that the model operates at 25 Hz. We kept the Gated Recurrent Unit (GRU) which the original implementation used to integrate latent states over a time window of 3 seconds, and used the transformer branches used to perform predictions from the GRU output for each of 15 upcoming samples, reaching .6 seconds into the future. We trained this model for 50 epochs on 360 hours of clean speech from the librispeech dataset (Panayotov et al., 2015), using a linear ramp to warm up the learning rate to the default value of .0002 over 10 epochs.

We propagated the audiobook stimulus through the network and stored the 256-dimensional outputs of the convolutional encoder, the GRU and each of the 15 transformer branches. We kept the first 32 PCA components for further modeling, which accounted for an average of 96.81% of variance, ranging from a minimum of 82.25% (GRU) to a maximum of 99.79% (Transformer at second forecast horizon).

##### 2.3.1.2 Wav2Vec 2.0

Secondly, we considered wav2vec 2.0 (Baevski et al., 2020), a further development of wav2vec (Schneider et al., 2019). The model consists of an encoder module using multiple layers of 1D convolutions to convert frames of the speech waveform of 25 ms duration into latent representations. These are then fed into 25 layers of transformer encoders that contextualise each frame of the latent representations with surrounding information. The latent representations from the convolutional encoder are quantised. A portion of quantised representations is then set to a masked token. The loss function is set up as a contrastive loss, where the model needs to maximise the similarity between the contextualised masked token and the true quantised latent representation prior to masking, and minimise the similarity between the contextualised masked token and random “negative” quantised latent representations. This effectively encourages the model to learn robust contextual dependencies. We used a large model pretrained and fine-tuned on 960 hours of librispeech as available from huggingface (*Huggingface.co*, n.d.).

We propagated the audiobook through the network and stored 1024-dimensional outputs of all 25 transformer layers at the native sampling rate of 50 Hz. We subsequently temporally downsampled it to 25 Hz and used the first 32 PCA components for our modeling purposes.

This number of components could account for an average of 73.97% of variance across layers, ranging from a minimum of 58.88% (layer 20) to a maximum of 99.90% (layer 25).

#### 2.3.2 Envelope prediction

Thirdly, we were interested in developing an autoregressive model akin to CPC, but restricted to the forecasting of sound energy fluctuations. The central idea of this network was thus to predict the speech envelope in the upcoming future from a segment of raw speech waveform. We were further interested in distributional predictions that would provide us with estimates of mean and variance parameterising a Gaussian probability density function.

We implemented this as a combination of a convolutional encoder paired with a gated recurrent unit (GRU) and a multilayer perceptron (MLP). The convolutional encoder used 7 layers of 1D convolutions, yielding a temporal downsampling factor of 640 and thereby going from 16 kHz in the speech input to 25 Hz of encoder outputs. Since we were interested in predicting absolute values of speech energy, we did not use batch- or layer-normalisation. A GRU with 3 layers of 100 units each integrated 50 convolutional encoder outputs over a time window of 2 seconds. Its output was fed into 3 fully connected layers with Gaussian Error Linear Unit (GELU, Hendrycks & Gimpel, 2016) nonlinearities. This was then weighted by linear readouts going through another GELU for a prediction of the mean *μ* and a softplus activation for a prediction of the standard deviation *σ*. We used 15 of such parameterisations of Gaussians to provide distributional predictions for 15 forecast horizons, reaching from .04 to .6 seconds into the future.

To define our loss, we extracted the speech envelope of the speech waveform immediately following the input to the network. We did this using a log-mel spectrogram as implemented in torchaudio with parameters following the GBFB toolbox (Schädler et al., 2012), using 31 mel-spaced channels covering a minimum frequency of 64 and a maximum frequency of 8001 Hz, 400 fast Fourier transform points, and a hop length of 160. We computed the decadic logarithm of the output values, multiplied the result by 20 and added a constant of 100. Finally we thresholded the output to a minimum of -20. We computed the mean of this spectrogram over channels and provided the output *y* at corresponding forecast horizons to a Gaussian negative log likelihood cost function:

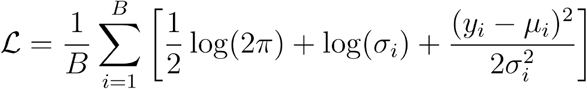

We trained the network for 50 epochs using a batch size *B* of 16, the adam optimiser (Kingma & Ba, 2014) with a learning rate of 1e-4 and a weight decay of 1e-6 on 360 hours of clean speech from the librispeech dataset (Panayotov et al., 2015). Additionally, we used a modification of the network where the speech input was first treated with the same denoising model described in section 2.1 prior to being propagated through the network. To ensure replicability (Mehrer et al., 2020), we repeated the training procedure 10 times for each of the two versions, each time with different random seeds.

We propagated the stimulus through each network at a sampling rate of 16 kHz and obtained dynamic predictions of mean and standard deviation. We downsampled these to 25 Hz for our encoding modeling purposes.

## 3 Two-stage analytical approach

Our analytical approach was based on a two-stage procedure. First, we were interested in assessing whether MEG power time courses were systematically related to the speech stimulus, and if so, what linear combination of the MEG sensors and frequencies would extract a suitable response component. Secondly, we were interested in comparing a range of feature spaces extracted from the speech stimulus with respect to their predictive power about the neuronal response component extracted in the first step.

All corresponding operations were organised within a nested cross-validation. Since the data was recorded from individual participants in 6 blocks of identical length, this was done with 6 outer folds. In each fold, we used 1 block as the test set and the remaining 5 blocks as the development set. In each of 5 inner folds, we further split the development set into 4 blocks of training and 1 block of validation data. This allowed us to optimise hyperparameters and minimise overfitting.

### 3.1.1 Canonical correlation analysis to extract a response component

We implemented the first stage as a Tikhonov regularised canonical component analysis (CCA). CCA finds a set of linear weights *W*_*enc*_ and *W*_*dec*_ that project two matrices *X* and *Y* onto loadings that are maximally correlated, where the resulting projections from each matrix are orthogonal to one another (Stratos, 2020). We assign a matrix consisting of the speech envelope as a column vector concatenated with delayed versions of itself to *X*, where the delays range from *t*_*min*_ to *t*_*max*_. Further, we assign a matrix of sensor level MEG power time courses to *Y*. The weights *W*_*enc*_ and *W*_*dec*_ thus correspond to paired encoding models (or multivariate temporal response functions, mTRFs, in essence finite impulse response filters, FIRs) and decoding models (or spatio-spectral filters), respectively. This can be implemented by first computing the whitened cross-covariance matrix of *X* and *Y*, using data from the training set:

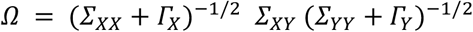

Here, *∑*_*XX*_, *∑*_*YY*_ and *∑*_*XY*_ are auto- and cross-covariance terms, respectively. We regularise the CCA by adding *Γ*_*X*_ and *Γ*_*Y*_ to the auto-covariance terms. To encourage smoothness across temporally neighbouring delays of the encoding model, we implement a Tikhonov regularisation with a discrete Laplace operator (Goutte et al., 2000), squared to a five-point stencil:

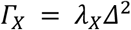

Where *Δ* is:

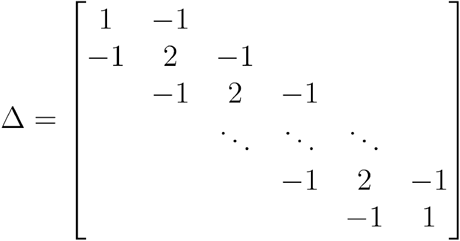

We use plain L2 regularisation for the decoding model:

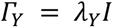

Where *I* is the identity matrix.

We can now obtain the left and right canonical directions *U* and *V* from singular value decomposition (SVD), using Matlab’s economy-sized decomposition:

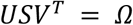

Lastly, we can obtain *W*_*enc*_ and *W*_*dec*_ by mapping the whitened canonical directions back to the original variable scales:

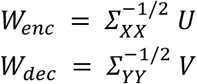

Data from the yet unseen validation set can now be transformed into the projections *X*_*proj*_ and *Y*_*proj*_ by multiplication:

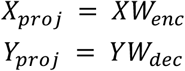

We can then use the negative Pearson correlation between *X*_*proj*_ and *Y*_*proj*_ as a loss function to continuously optimise the hyperparameters *t*_*min*_, *t*_*max*_, *λ*_*X*_ and *λ*_*Y*_. We find that we obtain the most stable solutions when optimising these hyperparameters only for the first canonical correlation. To do so, we use Bayesian adaptive direct search (BADS, Acerbi & Ma, 2017), a Gaussian process based black box optimisation tool, which we run with a maximum number of iterations of 200 multiplied by the number of hyperparameters and a maximum number of function evaluations of 500 multiplied by the number of hyperparameters. Additionally, we repeat this procedure 10 times, the first time with initial values as specified in Table S1, the following 9 times with initial values sampled from a uniform random distribution between the hyperparameter boundaries specified in Table S1. Across the 10 repetitions, we keep the hyperparameter combinations resulting in the best validation performance. Additionally, we apply *W*_*dec*_ to the training set portions of *Y* and store these projections for the second stage. Once this has been done for the 5 inner folds of the nested cross-validation, we choose that of the 5 combinations which is closest to the median across the inner folds. We now refit the CCA using the entire development set and measure the test set performance by applying the resulting *W*_*enc*_ and *W*_*dec*_ to portions of *X* and *Y* from the unseen fold. We also apply *W*_*dec*_ to the development set portions of *Y* and store these projections for the second stage.

To investigate which sensors and frequencies contributed to our component, we computed “pattern matrices” *P* (Haufe et al., 2014; Parra et al., 2005; van Vliet & Salmelin, 2020):

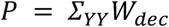

### 3.1.2 Source localisation of response component

To estimate where in the brain the response component extracted using CCA was generated, we used a linear model approach (Parra et al., 2005) predicting the canonical response component time courses from activity reconstructed for a range of grid points in source space. To do so, we used volume conductor models based on individual T1-weighted anatomical magnetic resonance imaging (MRI) scans which were aligned with the digitised head shapes using the iterative closest point algorithm. MRI scans were segmented to generate corrected-sphere volume conductor models (Nolte, 2003). We generated a grid of points in Montreal Neurological Institute (MNI) space at a resolution of 5 mm. We then transformed individual anatomies into MNI space and applied the inverses of these transformations to the grid in MNI space. This allowed us to estimate activity at locations distributed in individual spaces and visualise it in a common template space. We computed leadfields for these grids.

We next filtered the sensor level activity using the same bank of filters as in 1.2 and computed covariance matrices from the resulting filtered time domain data. We then constructed linearly constrained minimum variance beamformer spatial filters (Van Veen et al., 1997), using 5% regularisation and keeping all three orientations, separately for each block and frequency. We projected the spectrally filtered sensor level activity through these spatial filters and obtained the absolute values of the Hilbert transforms. Subsequently, we used ridge regression in a nested 6-fold cross-validation to predict the CCA response component from a matrix of (time x [frequency * orientation]) 82500 x [20 * 3], separately for each grid point. The split between training, validation and test sets was equal to that described in 3. To optimise the amount of L2 regularisation, we again used BADS, here with a maximum of iterations set to 1000, an initial *λ* of 2^17^ and boundaries ranging from 2^− 50^ to 2^50^. Essentially, this resulted in maps of Pearson correlation for each grid point, which we averaged across blocks and participants to visualise a group level source localisation.

### 3.2.1 Linear regression for feature space comparison

We implemented the second stage as a Tikhonov regularised linear regression predicting *Y*_*proj*_ (as obtained in the first stage) from various feature spaces extracted from the stimulus material. In essence, this is the same as obtaining *W*_*enc*_ from CCA for different stimulus feature spaces while keeping the decoder fixed, allowing direct comparisons of test set performances. To avoid leakage between training and testing sets within the same cross-validation scheme as in the first stage, we used different weights to obtain the predictee for inner and outer folds: We used the stored *Y*_*proj*_ obtained from *W*_*dec*_ fit to the training set of the respective CCA inner fold as the predictee in the inner folds of the second stage. We then measured the test set performance of the linear regression using the stored *Y*_*proj*_ obtained from *W*_*dec*_ trained on the development set of the respective outer fold as the predictee.

We obtain the regression weights by solving the normal equation of regularised linear regression:

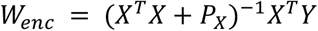

Here, *P*_*X*_ is defined in the same way as in the first stage, except for two modifications when *X* consists of multiple dimensions and/or subspaces. For multivariate feature spaces, we arrange *Δ* such that the discrete Laplace operator correctly addresses temporally neighbouring delays. For multiple subspaces, we provide individual *λ*_*X*_ for each subspace (Daube et al., 2019). We again used BADS to optimise the hyperparameters (*t*_*min*_, *t*_*max*_, *λ*_*X*_ per feature subspace) with the same numbers of maximum iterations and function evaluations as in the first stage, and the initial values and boundaries for hyperparameters specified in Table S2. We find that this hyperparameter optimisation needs to optimise the same objective function as in the first stage (Pearson correlation, not mean squared error) in order to achieve the same performance of *W*_*enc*_ obtained from CCA or regression with the speech envelope as predictor.

### 3.2.2 Phase response regularisation

During generalisation testing (see below), we found that the phase response of our *W*_*enc*_ as identified in the second stage of our analysis had a high variance especially for higher frequencies. We therefore devised a method to constrain the phase response. We did this with a modification of the loss function optimised using BADS in the second stage.

The default loss function is defined as:

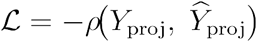

Where *ρ* denotes the Pearson correlation. The modified loss function is defined as:

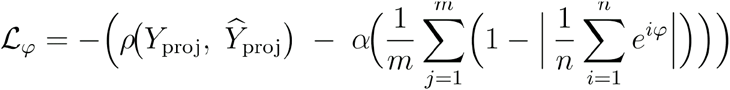

Here, *α* is a constant set to .01, *m* is the number of features in the respective feature space, *n* denotes the number of frequencies for which the phase response is computed and *φ* is the phase response of a given feature:

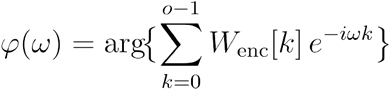

Where *o* is the number of delays of the filter *W*_*enc*_ and *ω* represents angular frequency in radians at which the phase response is sampled. We implement this using Matlab’s “phasez” function with its default number of 512 points.

This penalty effectively encourages a high mean resultant vector length across frequencies, or in other words, a constant phase response of the encoding filter.

## 4 Generalisation testing on controlled experiment

In addition to testing our encoding models on the passive audiobook listening MEG data, we were also interested in assessing how the model would generalise to an experimentally controlled condition (out-of-distribution relative to the speech stimulus material) known in the literature. Fujioka et al. (2012) describe an experiment in which participants listen to isochronous tones and report beta power modulations with respect to the tone onsets. In figure 3 of the original publication, grand average traces for each condition are provided alongside overlaid individual participant data, suggesting a high degree of between-participant consistency. We concluded that the grand average traces are a good representation even of single participant data and used a python tool (*Plotdigitizer*, n.d.) to convert grand average beta power modulations in response to the four experimental conditions into spreadsheets. This returned 113, 109, 104 and 123 ordered pairs of time in ms and beta power modulations, which we interpolated onto a common set of query points at 25 Hz, covering -.1 to .9 seconds of peri-onset time.

Next, we followed the original methods section to resynthesise the stimulus material used in the experiment. Specifically, we created a sine tone at 262 Hz lasting .06 seconds, which we multiplied with a hanning taper. We concatenated these tones to form trains of 15 seconds, spaced by stimulus onset asynchronies of either .78, .585 or .39 seconds for the three isochronous conditions, and spaced by a duration drawn from a uniform distribution *U*_[.39*s*.78*s*]_ for each of 50 random repetitions of the random condition. We propagated these stimuli through each of our acoustic feature spaces and networks and multiplied the output with the weights of the encoding models from section 3.2.1, separately for each fold of the cross-validation. We then epoched the predicted responses according to the onsets of the tones and correlated the trial averages with the data extracted from the figure 3 of the original publication (Fujioka et al., 2012). Finally, we averaged the correlations from the 50 repetitions of the random condition to obtain 6 Pearson correlations for each of four conditions per participant and model.

## 5 Statistical evaluations

We were interested in comparing the performance of sets of our models in various contexts. These performances were measured as Pearson correlations between observed and predicted time series, and were available from multiple folds for each participant. For the generalisation testing, these were available for each of 4 conditions. Prior to linear modeling, we Fisher-Z transformed the correlations, and then applied the inverse transform to samples from the posterior prior to visualisation. We used a Bayesian linear modeling approach as offered by the brms toolbox (Bürkner, 2017) to describe the structure of the respective set of correlations. To do so, we used weakly informative prior distributions that got easily overwhelmed by the likelihood, and drew a total of *N*_*s*_ *=* 16000 samples distributed across multiple chains, where each chain was warmed up with *N*_*w*_ *=* 1000 samples.

### 5.1 Comparison of acoustic and linguistic encoders for passive speech listening data

To model the comparison of the performances of encoding models when predicting outputs of the CCA decoder and the LCMV spatial filter, we used the following structure for each observation *i* of each participant *j*, predicting transformed correlations *z* from a binary vector *F* denoting the respective method of linearly combining sensors, using a weight vector for main effects *β* as well as participant-specific random intercepts *α* and random slopes *γ*

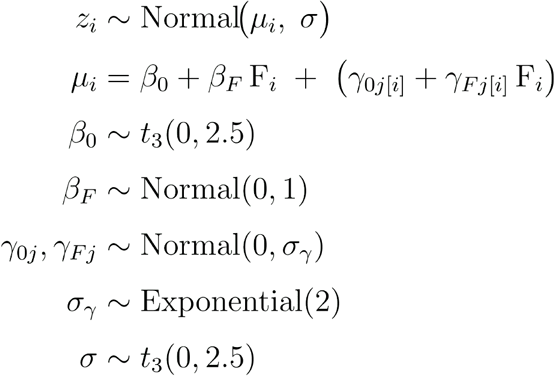

We used the same formula to model the comparisons of performances of acoustic and linguistic predictors on the passive speech listening data. In this case, *F* was a binary matrix denoting the feature space whose performance was measured.

### 5.2 Evaluation of generalisation testing

For the generalisation testing on the isochronous tones stimuli, we followed the same approach, but included three modifications. Firstly, we found a Skew Normal distribution to better match the data. Secondly, we expanded the model to a distributional model with a linear predictor not just for the location, but also for the scale and the skew. We express the corresponding parameters with *τ* and κ respectively. Thirdly, we included main effects for the four conditions as well as main effects for each combination of conditions and feature spaces. This model thus also used a binary matrix *C* denoting the four conditions of the experiment by Fujioka et al. (2012), and we use ⊗ to denote the row-wise Kronecker product between *C* and F. We found these choices to be supported by model comparisons including Gaussian, Student’s *t* and Skew Normal families with distributional and non-distributional options each. Our final choice featured the highest expected log pointwise predicted density.

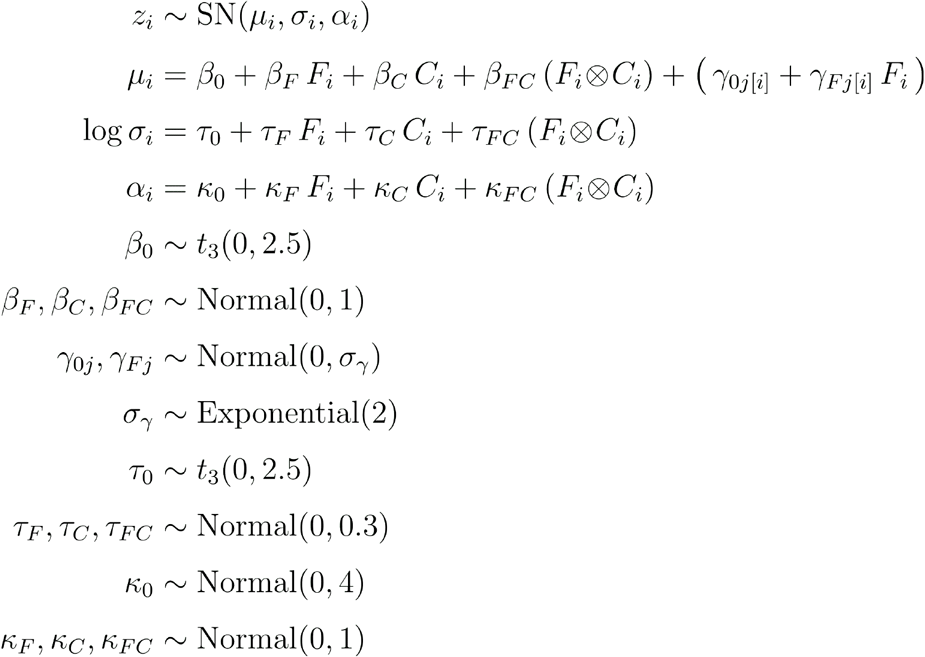

### 5.3 Comparison of encoders with and without phase regularisation

To evaluate the effect of the phase regularisation on encoding and generalisation performance, we fitted additional models using performances from encoding models using only the acoustic predictors but trained both without and with the phase regularisation. For models describing results of the performance in the speech listening data, we included a main effect and a random slope denoting the phase regularisation *P*. For models describing results of the generalisation testing, we additionally included a main effect denoting combinations of phase regularisation and the four experimental conditions, and did so for all three linear predictors of location, scale and skew. This resulted in the following formula for comparisons on the speech listening data:

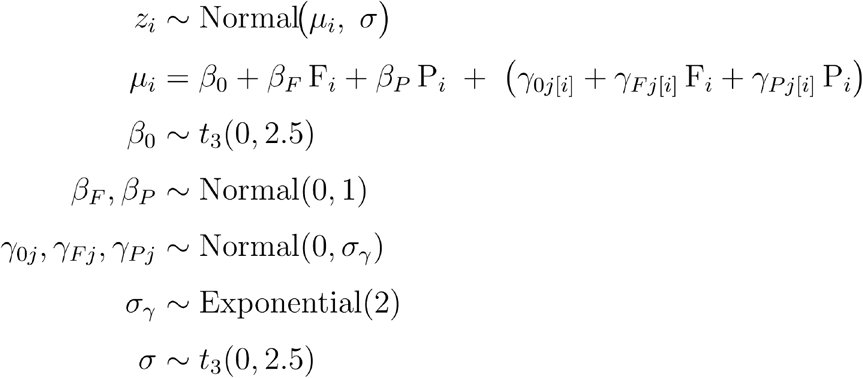

Further, we used the following formula for comparisons on the Fujioka generalisation testing:

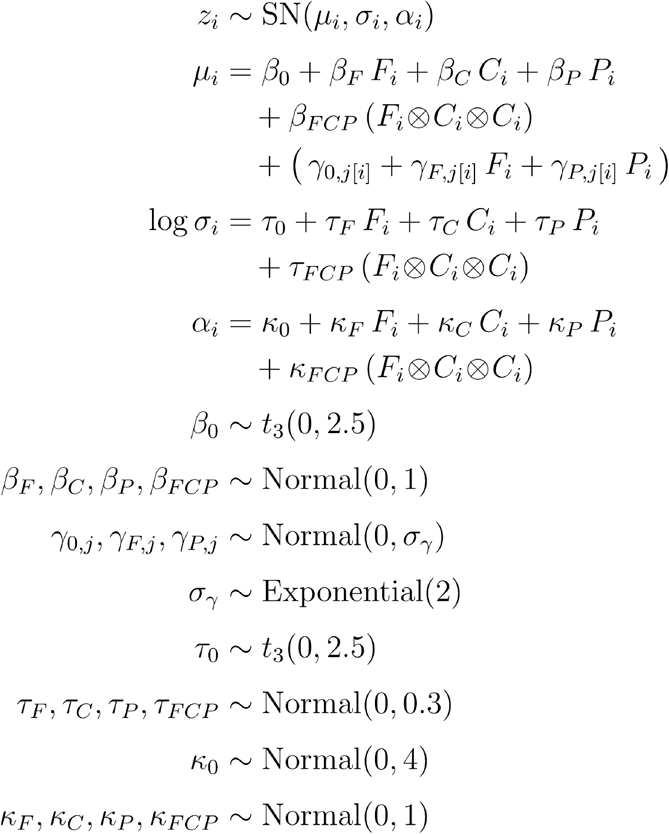

### 5.4 Comparison of effects for different models

Once the models were fit, we computed draws of the expected value of the posterior predictive distribution for hypothesis testing and visualisation. To do so, we constructed counterfactual design matrices isolating individual predictor combinations of interest and ran our models with the posterior draws from all parameters. This took the following form for comparisons on the speech listening data and the generalisation testing:

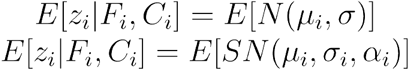

And the following adapted form for the evaluation of the phase regularisation:

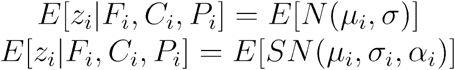

This procedure yielded, for each model, a matrix of draws from the expected value of the posterior predictive distribution where each column represented the conditional expectation for a specific predictor combination (e.g. a single feature space). To evaluate directed hypotheses, e.g. between any two feature spaces *A* and *B*, we computed corresponding differences directly from this matrix and obtained the fraction of samples in favour of a given hypothesis as:

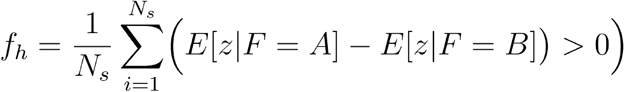

This ratio can be used to form an “evidence ratio” 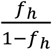, which corresponds to a Bayes Factor computed via the Savage-Dickey density ratio method. According to Jeffreys (1998), Bayes Factors can be mapped onto a scale of interpretation ranging from “barely worth mentioning” ([10^0^, 10^.5^]) to “substantial” ([10^.5^, 10^1^]), “strong” ([10^1^, 10^1.5^]), “very strong” ([10^1.5^, 10^2^]) and “decisive” (> 10^2^). This thus maps onto the following intervals of *f*_*h*_: “barely worth mentioning” ([.5000, .7597]) to “substantial” ([.7597, .9091]), “strong” ([.9091, .9693]), “very strong” ([.9693, .9901]) and “decisive” (> .9901).

## Supplemental material

**Figure S1:**
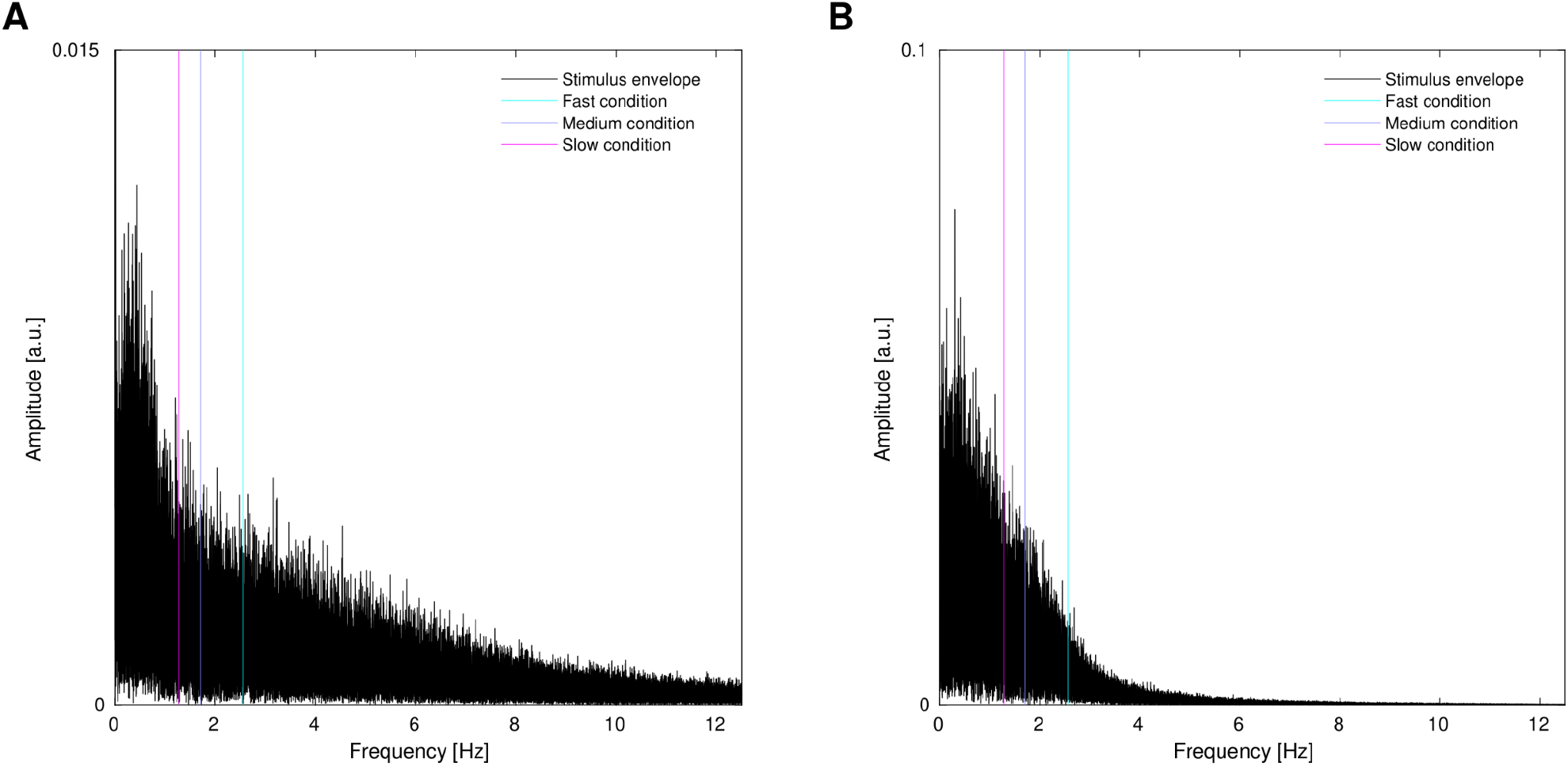
Less signal at frequencies corresponding to faster conditions provides less constraint for phase responses **A**) Modulation power spectrum of the speech stimulus (i.e. power spectrum of the speech envelope). The frequencies corresponding to the three isochronous conditions from Fujioka et al. (2012) are overlaid as vertical lines. **B**) Modulation power spectrum of the neural response component (i.e. power spectrum of power time course) of an exemplary participant. The frequencies corresponding to the three isochronous conditions from Fujioka et al. (2012) are overlaid as vertical lines. While there is a relatively constant amount of energy in the modulation power spectrum of the speech signal (A), the beta power responses have less than half of the energy at a frequency corresponding to the fastest condition compared to the slowest condition.

**Figure S2:**
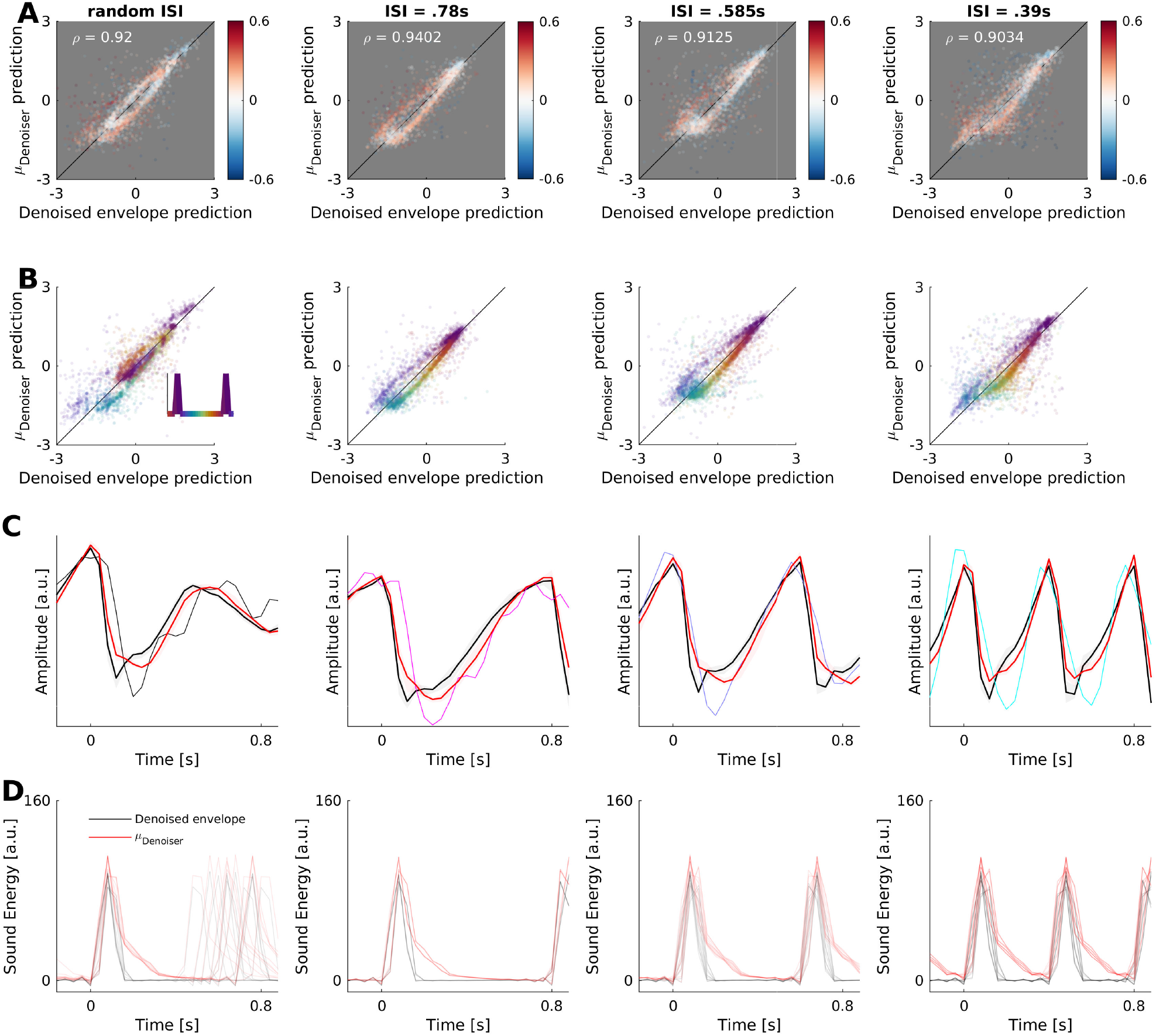
Evaluations of predictions from acoustic features and network features on all four conditions from Fujioka et al. (2012). **A**) Scatter plots of predictions from acoustic feature (denoised envelope) and network feature (forecasted mean in network predicting denoised envelope). Each data point is a single predicted time point of a single fold and participant. Data points are coloured according to differences of absolute residuals, where positive values denote smaller residuals (i.e. better predictions) for the network feature. Scale is third order symmetric logarithm. **B**) Same data as in A, but coloured according to time relative to tone onset. Inset in leftmost panel shows colours mapped on tone triggers over time. **C**) Beta power dynamics at tone onset, showing observations made by Fujioka et al. (2012) as well as predictions from denoised envelope and network feature (mean ± 95% bootstrapped confidence interval across 24 participants). **D**) Comparison of raw acoustic features and network features (i.e. the inputs to TRF modeling). Each line is a single trial. All columns in A, B, C and D show (from left to right) random, slow (.78 s), medium (.585 s) and fast (.39 s) conditions.

**Table S1:** Initial values and boundaries for hyperparameters in BADS optimisation of regularised CCA.

| hyperparameter | Initial value | Lower boundary | Upper boundary |
| --- | --- | --- | --- |
| $t_{min}$ [s] | -1.0 | -3.0 | 0.4 |
| $t_{max}$ [s] | 1.0 | -0.4 | 3.0 |
| $\log_2(\lambda_X)$ | 0 | -50 | 50 |
| $\log_2(\lambda_Y)$ | 0 | -50 | 50 |

**Table S2:** Initial values and boundaries for hyperparameters in BADS optimisation of regularised encoding models.

| hyperparameter | Initial value | Lower boundary | Upper boundary |
| --- | --- | --- | --- |
| $t_{min}$ | -0.2 | -1.5 | 0.48 |
| $t_{max}$ | 0.8 | 0.2 | 2.48 |
| $\log_2(\lambda_X)$ | 19 | -30 | 30 |

## References

Acerbi, L., & Ma, W. J. (2017). Practical Bayesian Optimization for Model Fitting with Bayesian Adaptive Direct Search. Advances in Neural Information Processing Systems, 30.

Ahissar, E., Nagarajan, S., Ahissar, M., Protopapas, A., Mahncke, H., & Merzenich, M. M. (2001). Speech comprehension is correlated with temporal response patterns recorded from auditory cortex. Proceedings of the National Academy of Sciences of the United States of America, 98(23), 13367–13372.

Anderson, A. J., Davis, C., & Lalor, E. C. (2024). Deep-learning models reveal how context and listener attention shape electrophysiological correlates of speech-to-language transformation. PLoS Computational Biology, 20(11), e1012537.

Antonello, R., & Huth, A. (2024). Predictive coding or just feature discovery? An alternative account of why language models fit brain data. Neurobiology of Language (Cambridge, Mass.), 5(1), 64–79.

Armeni, K., Willems, R. M., van den Bosch, A., & Schoffelen, J.-M. (2019). Frequency-specific brain dynamics related to prediction during language comprehension. NeuroImage, 198, 283–295.

Arnal, L. H., Doelling, K. B., & Poeppel, D. (2015). Delta-beta coupled oscillations underlie temporal prediction accuracy. Cerebral Cortex (New York, N.Y.: 1991), 25(9), 3077–3085.

Baevski, A., Zhou, H., Mohamed, A., & Auli, M. (2020). wav2vec 2.0: a framework for self-supervised learning of speech representations. Proceedings of the 34th International Conference on Neural Information Processing Systems, Article Article 1044.

Betti, V., Della Penna, S., de Pasquale, F., & Corbetta, M. (2021). Spontaneous beta band rhythms in the predictive coding of natural stimuli. The Neuroscientist, 27(2), 184–201.

Bilenko, N. Y., & Gallant, J. L. (2016). Pyrcca: Regularized kernel canonical correlation analysis in Python and its applications to neuroimaging. Frontiers in Neuroinformatics, 10, 49.

Boersma, P. (2001). PRAAT, a system for doing phonetics by computer. Glot International, 5, 341–345.

Bowers, J. S., Malhotra, G., Dujmovic, M., Llera Montero, M., Tsvetkov, C., Biscione, V., Puebla, G., Adolfi, F., Hummel, J. E., Heaton, R. F., Evans, B. D., Mitchell, J., & Blything, R. (2022). Deep problems with neural network models of human vision. The Behavioral and Brain Sciences, 46, e385.

Brodbeck, C., Hannagan, T., & Magnuson, J. S. (2025). Recurrent neural networks as neuro-computational models of human speech recognition. PLoS Computational Biology, 21(7), e1013244.

Brodbeck, C., & Simon, J. Z. (2020). Continuous speech processing. 18, 25–31.

Bürkner, P.-C. (2017). Brms: An R package for Bayesian multilevel models using Stan. Journal of Statistical Software, 80(1), 1–28.

Buzsáki, G., & Draguhn, A. (2004). Neuronal oscillations in cortical networks. Science (New York, N.Y.), 304(5679), 1926–1929.

Carvalho, W., & Lampinen, A. (2025). Naturalistic Computational Cognitive Science: Towards generalizable models and theories that capture the full range of natural behavior. In arXiv [q-bio.NC]. arXiv. 10.48550/arXiv.2502.20349

Chalas, N., Daube, C., Kluger, D. S., Abbasi, O., Nitsch, R., & Gross, J. (2022). Multivariate analysis of speech envelope tracking reveals coupling beyond auditory cortex. NeuroImage, 258(119395), 119395.

Chalas, N., Daube, C., Kluger, D. S., Abbasi, O., Nitsch, R., & Gross, J. (2023). Speech onsets and sustained speech contribute differentially to delta and theta speech tracking in auditory cortex. Cerebral Cortex (New York, N.Y.: 1991), 33(10), 6273–6281.

Conwell, C., Prince, J. S., Kay, K. N., Alvarez, G. A., & Konkle, T. (2024). A large-scale examination of inductive biases shaping high-level visual representation in brains and machines. Nature Communications, 15(1), 9383.

Crosse, M. J., Di Liberto, G. M., Bednar, A., & Lalor, E. C. (2016). The multivariate temporal response function (mTRF) toolbox: A MATLAB toolbox for relating neural signals to continuous stimuli. Frontiers in Human Neuroscience, 10, 604.

Daube, C., Gross, J., & Ince, R. A. A. (2022). A whitening approach for Transfer Entropy permits the application to narrow-band signals. In arXiv [q-bio.NC]. arXiv. http://arxiv.org/abs/2201.02461

Daube, C., Ince, R. A. A., & Gross, J. (2019). Simple acoustic features can explain phoneme-based predictions of cortical responses to speech. Current Biology: CB, 29(12), 1924– 1937.e9.

Daube, C., Xu, T., Zhan, J., Webb, A., Ince, R. A. A., Garrod, O. G. B., & Schyns, P. G. (2021). Grounding deep neural network predictions of human categorization behavior in understandable functional features: The case of face identity. Patterns (New York, N.Y.), 2(10), 100348.

David, S. V., Vinje, W. E., & Gallant, J. L. (2004). Natural stimulus statistics alter the receptive field structure of v1 neurons. The Journal of Neuroscience: The Official Journal of the Society for Neuroscience, 24(31), 6991–7006.

de Cheveigné, A., & Nelken, I. (2019). Filters: When, why, and how (not) to use them. Neuron, 102(2), 280–293.

de Cheveigné, A., Wong, D. D. E., Di Liberto, G. M., Hjortkjær, J., Slaney, M., & Lalor, E. (2018). Decoding the auditory brain with canonical component analysis. NeuroImage, 172, 206–216.

Défossez, A., Synnaeve, G., & Adi, Y. (2020, October 25). Real time speech enhancement in the waveform domain. Interspeech 2020. Interspeech 2020. 10.21437/interspeech.2020-2409

de Lange, F. P., Rahnev, D. A., Donner, T. H., & Lau, H. (2013). Prestimulus oscillatory activity over motor cortex reflects perceptual expectations. The Journal of Neuroscience: The Official Journal of the Society for Neuroscience, 33(4), 1400–1410.

Di Liberto, G. M., O’Sullivan, J. A., & Lalor, E. C. (2015). Low-frequency cortical entrainment to speech reflects phoneme-level processing. Current Biology: CB, 25(19), 2457–2465.

Ding, N., Melloni, L., Zhang, H., Tian, X., & Poeppel, D. (2016). Cortical tracking of hierarchical linguistic structures in connected speech. Nature Neuroscience, 19(1), 158–164.

Ding, N., & Simon, J. Z. (2012). Emergence of neural encoding of auditory objects while listening to competing speakers. Proceedings of the National Academy of Sciences of the United States of America, 109(29), 11854–11859.

Dmochowski, J. P., Ki, J. J., DeGuzman, P., Sajda, P., & Parra, L. C. (2018). Extracting multidimensional stimulus-response correlations using hybrid encoding-decoding of neural activity. NeuroImage, 180(Pt A), 134–146.

Doelling, K. B., Assaneo, M. F., Bevilacqua, D., Pesaran, B., & Poeppel, D. (2019). An oscillator model better predicts cortical entrainment to music. Proceedings of the National Academy of Sciences of the United States of America, 116(20), 10113–10121.

Doerig, A., Sommers, R. P., Seeliger, K., Richards, B., Ismael, J., Lindsay, G. W., Kording, K. P., Konkle, T., van Gerven, M. A. J., Kriegeskorte, N., & Kietzmann, T. C. (2023). The neuroconnectionist research programme. Nature Reviews. Neuroscience, 24(7), 431–450.

Donhauser, P. W., & Baillet, S. (2020). Two distinct neural timescales for predictive speech processing. Neuron, 105(2), 385–393.e9.

Edelman, G. M., & Gally, J. A. (2001). Degeneracy and complexity in biological systems. Proceedings of the National Academy of Sciences of the United States of America, 98(24), 13763–13768.

Elmoznino, E., & Bonner, M. F. (2024). High-performing neural network models of visual cortex benefit from high latent dimensionality. PLoS Computational Biology, 20(1), e1011792.

Feather, J., Leclerc, G., Madry, A., & McDermott, J. H. (2023). Model metamers reveal divergent invariances between biological and artificial neural networks. Nature Neuroscience, 26(11), 2017–2034.

Fujioka, T., Trainor, L. J., Large, E. W., & Ross, B. (2012). Internalized timing of isochronous sounds is represented in neuromagnetic β oscillations. The Journal of Neuroscience: The Official Journal of the Society for Neuroscience, 32(5), 1791–1802.

Giordano, B. L., Esposito, M., Valente, G., & Formisano, E. (2023). Intermediate acoustic-to-semantic representations link behavioral and neural responses to natural sounds. Nature Neuroscience, 26(4), 664–672.

Giroud, J., Trébuchon, A., Mercier, M., Davis, M. H., & Morillon, B. (2024). The human auditory cortex concurrently tracks syllabic and phonemic timescales via acoustic spectral flux. Science Advances, 10(51), eado8915.

Golan, T., Guo, W., Schütt, H. H., & Kriegeskorte, N. (2022). Distinguishing representational geometries with controversial stimuli: Bayesian experimental design and its application to face dissimilarity judgments. In arXiv [q-bio.NC]. arXiv. http://arxiv.org/abs/2211.15053

Golan, T., Raju, P. C., & Kriegeskorte, N. (2020). Controversial stimuli: Pitting neural networks against each other as models of human cognition. Proceedings of the National Academy of Sciences of the United States of America, 117(47), 29330–29337.

Goutte, C., Nielsen, F. A., & Hansen, L. K. (2000). Modeling the haemodynamic response in fMRI using smooth FIR filters. IEEE Transactions on Medical Imaging, 19(12), 1188–1201.

Grabenhorst, M., Poeppel, D., & Michalareas, G. (2025). Neural signatures of temporal anticipation in human cortex represent event probability density. Nature Communications, 16(1), 2602.

Guest, O., & Martin, A. E. (2023). On logical inference over brains, behaviour, and artificial neural networks. Computational Brain & Behavior, 6(2), 213–227.

Gwilliams, L., & King, J.-R. (2020). Recurrent processes support a cascade of hierarchical decisions. eLife, 9, e56603.

Gwilliams, L., Marantz, A., Poeppel, D., & King, J.-R. (2025). Hierarchical dynamic coding coordinates speech comprehension in the human brain. Proceedings of the National Academy of Sciences of the United States of America, 122(42), e2422097122.

Haegens, S., Nácher, V., Luna, R., Romo, R., & Jensen, O. (2011). α-Oscillations in the monkey sensorimotor network influence discrimination performance by rhythmical inhibition of neuronal spiking. Proceedings of the National Academy of Sciences of the United States of America, 108(48), 19377–19382.

Hamilton, L. S., & Huth, A. G. (2020). The revolution will not be controlled: natural stimuli in speech neuroscience. Language, Cognition and Neuroscience, 35(5), 573–582.

Haufe, S., Meinecke, F., Görgen, K., Dähne, S., Haynes, J.-D., Blankertz, B., & Bießmann, F. (2014). On the interpretation of weight vectors of linear models in multivariate neuroimaging. NeuroImage, 87, 96–110.

Heilbron, M., Armeni, K., Schoffelen, J.-M., Hagoort, P., & de Lange, F. P. (2022). A hierarchy of linguistic predictions during natural language comprehension. Proceedings of the National Academy of Sciences of the United States of America, 119(32), e2201968119.

Hendrycks, D., & Gimpel, K. (2016). Gaussian Error Linear Units (GELUs). In arXiv [cs.LG]. arXiv. http://arxiv.org/abs/1606.08415

Henry, M. J., & Herrmann, B. (2014). Low-frequency neural oscillations support dynamic attending in temporal context. Timing & Time Perception (Leiden, Netherlands), 2(1), 62– 86.

Herrmann, B., Henry, M. J., Haegens, S., & Obleser, J. (2016). Temporal expectations and neural amplitude fluctuations in auditory cortex interactively influence perception. NeuroImage, 124(Pt A), 487–497.

Hovsepyan, S., Olasagasti, I., & Giraud, A.-L. (2023). Rhythmic modulation of prediction errors: A top-down gating role for the beta-range in speech processing. PLoS Computational Biology, 19(11), e1011595. huggingface.co. (n.d.). Hugging Face. Retrieved October 16, 2025, from https://huggingface.co/facebook/wav2vec2-large-960h

Jeffreys, H. (1998). The theory of probability. Oxford University Press.

Keitel, A., Gross, J., & Kayser, C. (2018). Perceptually relevant speech tracking in auditory and motor cortex reflects distinct linguistic features. PLoS Biology, 16(3), e2004473.

Kell, A. J. E., Yamins, D. L. K., Shook, E. N., Norman-Haignere, S. V., & McDermott, J. H. (2018). A task-optimized neural network replicates human auditory behavior, predicts brain responses, and reveals a cortical processing hierarchy. Neuron, 98(3), 630–644.e16.

Khalighinejad, B., Herrero, J. L., Mehta, A. D., & Mesgarani, N. (2019). Adaptation of the human auditory cortex to changing background noise. Nature Communications, 10(1), 2509.

Kim, S.-G. (2022). On the encoding of natural music in computational models and human brains. Frontiers in Neuroscience, 16, 928841.

Kingma, D. P., & Ba, J. (2014). Adam: A method for stochastic optimization. In arXiv [cs.LG]. arXiv. http://arxiv.org/abs/1412.6980

Krakauer, J. W., Ghazanfar, A. A., Gomez-Marin, A., MacIver, M. A., & Poeppel, D. (2017). Neuroscience needs behavior: Correcting a reductionist bias. Neuron, 93(3), 480–490.

Kriegeskorte, N., & Douglas, P. K. (2018). Cognitive computational neuroscience. Nature Neuroscience, 21(9), 1148–1160.

Lampinen, A. K., Chan, S. C. Y., Li, Y., & Hermann, K. (2025). Representation biases: will we achieve complete understanding by analyzing representations? In arXiv [q-bio.NC]. arXiv. http://arxiv.org/abs/2507.22216

Leske, S., Endestad, T., Volehaugen, V., Foldal, M. D., Blenkmann, A. O., Solbakk, A.-K., & Danielsen, A. (2025). Beta oscillations predict the envelope sharpness in a rhythmic beat sequence. Scientific Reports, 15(1), 3510.

Liu, J., & Zhang, Y. (2017). In-order transition-based constituent parsing. Transactions of the Association for Computational Linguistics, 5, 413–424.

Marslen-Wilson, W., & Tyler, L. K. (1980). The temporal structure of spoken language understanding. Cognition, 8(1), 1–71.

Mehrer, J., Spoerer, C. J., Kriegeskorte, N., & Kietzmann, T. C. (2020). Individual differences among deep neural network models. Nature Communications, 11(1), 5725.

Mesgarani, N., David, S. V., Fritz, J. B., & Shamma, S. A. (2014). Mechanisms of noise robust representation of speech in primary auditory cortex. Proceedings of the National Academy of Sciences of the United States of America, 111(18), 6792–6797.

Miller, G. A., & Chomsky, N. (1963). Finitary models of language users. In D. Luce (Ed.), Handbook of Mathematical Psychology (pp. 2–419). John Wiley & Sons.

Miller, G. A., & Isard, S. (1964). Free recall of self-embedded english sentences. Information and Control, 7(3), 292–303.

Millet, J., Caucheteux, C., Orhan, P., Boubenec, Y., Gramfort, A., Dunbar, E., Pallier, C., & King, J.-R. (2022). Toward a realistic model of speech processing in the brain with self-supervised learning. In arXiv [q-bio.NC]. arXiv. http://arxiv.org/abs/2206.01685

Naselaris, T., Kay, K. N., Nishimoto, S., & Gallant, J. L. (2011). Encoding and decoding in fMRI. NeuroImage, 56(2), 400–410.

Niv, Y. (2021). The primacy of behavioral research for understanding the brain. Behavioral Neuroscience, 135(5), 601–609.

Nolte, G. (2003). The magnetic lead field theorem in the quasi-static approximation and its use for magnetoencephalography forward calculation in realistic volume conductors. Physics in Medicine and Biology, 48(22), 3637–3652.

Obleser, J. (2025). Metacognition in the listening brain. Trends in Neurosciences, 48(2), 100– 112.

Oganian, Y., & Chang, E. F. (2019). A speech envelope landmark for syllable encoding in human superior temporal gyrus. Science Advances, 5(11), eaay6279.

Oostenveld, R., Fries, P., Maris, E., & Schoffelen, J.-M. (2011). FieldTrip: Open source software for advanced analysis of MEG, EEG, and invasive electrophysiological data. Computational Intelligence and Neuroscience, 2011, 156869.

O’Sullivan, J. A., Power, A. J., Mesgarani, N., Rajaram, S., Foxe, J. J., Shinn-Cunningham, B. G., Slaney, M., Shamma, S. A., & Lalor, E. C. (2015). Attentional selection in a cocktail party environment can be decoded from single-trial EEG. Cerebral Cortex (New York, N.Y.: 1991), 25(7), 1697–1706.

Panayotov, V., Chen, G., Povey, D., & Khudanpur, S. (2015). Librispeech: An ASR corpus based on public domain audio books. 2015 IEEE International Conference on Acoustics, Speech and Signal Processing (ICASSP), 5206–5210.

Parra, L. C., Spence, C. D., Gerson, A. D., & Sajda, P. (2005). Recipes for the linear analysis of EEG. NeuroImage, 28(2), 326–341.

Pesnot Lerousseau, J., Trébuchon, A., Morillon, B., & Schön, D. (2021). Frequency selectivity of persistent cortical oscillatory responses to auditory rhythmic stimulation. The Journal of Neuroscience: The Official Journal of the Society for Neuroscience, 41(38), 7991–8006.

plotdigitizer. (n.d.). PyPI. Retrieved March 17, 2026, from https://pypi.org/project/plotdigitizer/

Poeppel, D., & Embick, D. (2005). Defining the relation between linguistics and neuroscience. In Cutler (Ed.), Twenty-first century psycholinguistics: Four cornerstones (pp. 103–120). Lawrence Erlbaum.

Putnam, H. (1967). Psychological predicates (W.H. Capitan & D.D. Merrill (Eds.); pp. 37–48). University of Pittsburgh Press.

Qi, P., Zhang, Y., Zhang, Y., Bolton, J., & Manning, C. D. (2020). Stanza: A python natural language processing toolkit for many human languages. In A. Celikyilmaz & T.-H. Wen (Eds.), Proceedings of the 58th Annual Meeting of the Association for Computational Linguistics: System Demonstrations (pp. 101–108). Association for Computational Linguistics.

Radford, A., Wu, J., Child, R., Luan, D., Amodei, D., Sutskever, I., & Others. (2019). Language models are unsupervised multitask learners. OpenAI Blog, 1(8), 9.

Rieke, F., Bodnar, D. A., & Bialek, W. (1995). Naturalistic stimuli increase the rate and efficiency of information transmission by primary auditory afferents. Proceedings. Biological Sciences, 262(1365), 259–265.

Riviere, M., Joulin, A., Mazare, P.-E., & Dupoux, E. (2020). Unsupervised pretraining transfers well across languages. ICASSP 2020 - 2020 IEEE International Conference on Acoustics, Speech and Signal Processing (ICASSP), 7414–7418.

Rust, N. C., & Movshon, J. A. (2005). In praise of artifice. Nature Neuroscience, 8(12), 1647– 1650.

Saleh, M., Reimer, J., Penn, R., Ojakangas, C. L., & Hatsopoulos, N. G. (2010). Fast and slow oscillations in human primary motor cortex predict oncoming behaviorally relevant cues. Neuron, 65(4), 461–471.

Schädler, M. R., Meyer, B. T., & Kollmeier, B. (2012). Spectro-temporal modulation subspace-spanning filter bank features for robust automatic speech recognition. The Journal of the Acoustical Society of America, 131(5), 4134–4151.

Schaeffer, R., Khona, M., Chandra, S., Ostrow, M., Miranda, B., & Koyejo, S. (2024, October 10). Position: Maximizing Neural Regression Scores May Not Identify Good Models of the Brain. UniReps: 2nd Edition of the Workshop on Unifying Representations in Neural Models. https://openreview.net/forum?id=vbtj05J68r

Schneider, S., Baevski, A., Collobert, R., & Auli, M. (2019, September 15). wav2vec: Unsupervised Pre-Training for Speech Recognition. Interspeech 2019. Interspeech 2019. 10.21437/interspeech.2019-1873

Schrimpf, M., Kubilius, J., Lee, M. J., Ratan Murty, N. A., Ajemian, R., & DiCarlo, J. J. (2020). Integrative benchmarking to advance neurally mechanistic models of human intelligence. Neuron, 108(3), 413–423.

Schroeder, C. E., & Lakatos, P. (2009). Low-frequency neuronal oscillations as instruments of sensory selection. Trends in Neurosciences, 32(1), 9–18.

Schyns, P. G., Snoek, L., & Daube, C. (2022). Degrees of algorithmic equivalence between the brain and its DNN models. Trends in Cognitive Sciences, 26(12), 1090–1102.

Schyns, P. G., Snoek, L., & Daube, C. (2023). Stimulus models test hypotheses in brains and DNNs. Trends in Cognitive Sciences, 27(3), 216–217.

Sedley, W., Gander, P. E., Kumar, S., Kovach, C. K., Oya, H., Kawasaki, H., Howard, M. A., & Griffiths, T. D. (2016). Neural signatures of perceptual inference. eLife, 5, e11476.

Speer, R., Chin, J., Lin, A., Nathan, L., & Jewett, S. (2016). wordfreq: v1.5.1. Zenodo. 10.5281/ZENODO.61937

Stratos, K. (2020). A Hitchhiker’s Guide to PCA and CCA. https://karlstratos.com/notes/pca_cca.pdf

Theunissen, F. E., Sen, K., & Doupe, A. J. (2000). Spectral-temporal receptive fields of nonlinear auditory neurons obtained using natural sounds. The Journal of Neuroscience: The Official Journal of the Society for Neuroscience, 20(6), 2315–2331.

Tukey, J. W. (1980). We need both exploratory and confirmatory. The American Statistician, 34(1), 23.

Vaidya, A. R., Jain, S., & Huth, A. G. (2022). Self-supervised models of audio effectively explain human cortical responses to speech. In arXiv [cs.CL]. arXiv. http://arxiv.org/abs/2205.14252

Vandecappelle, S., Deckers, L., Das, N., Ansari, A. H., Bertrand, A., & Francart, T. (2021). EEG-based detection of the locus of auditory attention with convolutional neural networks. eLife, 10(e56481), e56481.

van den Oord, A., Li, Y., & Vinyals, O. (2018). Representation learning with Contrastive Predictive Coding. In arXiv [cs.LG]. arXiv. http://arxiv.org/abs/1807.03748

VanRullen, R. (2017). Perception science in the age of deep neural networks. Frontiers in Psychology, 8, 142.

Van Veen, B. D., van Drongelen, W., Yuchtman, M., & Suzuki, A. (1997). Localization of brain electrical activity via linearly constrained minimum variance spatial filtering. IEEE Transactions on Bio-Medical Engineering, 44(9), 867–880.

van Vliet, M., & Salmelin, R. (2020). Post-hoc modification of linear models: Combining machine learning with domain information to make solid inferences from noisy data. NeuroImage, 204(116221), 116221.

Varoquaux, G., Raamana, P. R., Engemann, D. A., Hoyos-Idrobo, A., Schwartz, Y., & Thirion, (2017). Assessing and tuning brain decoders: Cross-validation, caveats, and guidelines. NeuroImage, 145(Pt B), 166–179.

Wang, H.-T., Smallwood, J., Mourao-Miranda, J., Xia, C. H., Satterthwaite, T. D., Bassett, D. S., & Bzdok, D. (2020). Finding the needle in a high-dimensional haystack: Canonical correlation analysis for neuroscientists. NeuroImage, 216(116745), 116745.

Weissbart, H., & Martin, A. E. (2024). The structure and statistics of language jointly shape cross-frequency neural dynamics during spoken language comprehension. Nature Communications, 15(1), 8850.

Wolf, T., Debut, L., Sanh, V., Chaumond, J., Delangue, C., Moi, A., Cistac, P., Rault, T., Louf, R., Funtowicz, M., Davison, J., Shleifer, S., von Platen, P., Ma, C., Jernite, Y., Plu, J., Xu, C., Le Scao, T., Gugger, S., … Rush, A. (2020). Transformers: State-of-the-art natural language processing. In Q. Liu & D. Schlangen (Eds.), Proceedings of the 2020 Conference on Empirical Methods in Natural Language Processing: System Demonstrations (pp. 38–45). Association for Computational Linguistics.

Yuan, J., & Liberman, M. (2008). Speaker identification on the SCOTUS corpus. The Journal of the Acoustical Society of America, 123(5_Supplement), 3878–3878.

Zioga, I., Weissbart, H., Lewis, A. G., Haegens, S., & Martin, A. E. (2023). Naturalistic spoken language comprehension is supported by alpha and beta oscillations. The Journal of Neuroscience: The Official Journal of the Society for Neuroscience, 43(20), 3718–3732.

Zou, J., Xu, C., Luo, C., Jin, P., Gao, J., Li, J., Gao, J., Ding, N., & Luo, B. (2021). Θ-band cortical tracking of the speech envelope shows the linear phase property. eNeuro, 8(4), ENEURO.0058–21.2021.

